# Landscape structural complexity influences the global distribution of ecosystems and people

**DOI:** 10.1101/2024.12.28.630608

**Authors:** James Cant, Nina Schiettekatte, Elizabeth M. P. Madin, Arie C. Seijmonsbergen, Kenneth F. Rijsdijk, Joshua S. Madin, Maria Dornelas

## Abstract

The three-dimensional structure of land- and seascapes links biotic and abiotic processes to shape how organisms use their environments. Much effort has been dedicated to understanding this relationship between structural complexity and biodiversity. However, tailored, system-specific approaches to quantifying structural complexity preclude cross-system comparisons. Expanding a geometric framework developed for quantifying the structural complexity of local-scale surface environments, we generate high-resolution, global maps of the key complexity measures of height range, rugosity, and fractal dimension. We illustrate how patterns in these complexity attributes correspond with the distribution of ecosystems, land cover types, and the presence of people across the globe. Using geometric tools to understand how structural complexity is associated with the arrangement of natural and socio-ecosystems, we can better predict their responses to ongoing global change.

## INTRODUCTION

The physical structure of natural environments sets the stage on which ecological, geological, and climatic processes shape global biodiversity and socio-ecosystems (*1*, *2*). Climatic regimes are a major determinant of global species richness trends (*3*). However, abiotic topography modulates the spatio-temporal configuration of climate systems, thereby mediating local climate exposure, resource availability, and resource access (*4–6*). Indeed, the contemporary arrangement of global biodiversity reflects how over time the interactions between topographic and climatic regimes have reconfigured local environments and the biotic communities they support (*7–11*). Some conservation approaches, such as *Conserving Nature’s Stage*, a global initiative aimed at incorporating landscape features in conservation planning (*12*), even argue that protecting the enduring physical structure of natural habitats is a more effective use of conservation resources than safeguarding more transitory biotic assemblages (*13–17*). The structure of natural environments also contributes to human wellbeing, provides key cultural benefits, and supports agricultural diversification (*2*, *18*). Because of this ecological and socioeconomic value, and the tightly interwoven relationship between global biological and geophysical processes, there is a critical need for better integration of topographic structure into our understanding of the mechanisms refining global biodiversity patterns (*19*). Identifying scalable metrics applicable across systems is thus essential for quantifying how abiotic structural complexity helps to shape ecological systems worldwide.

Establishing the association between abiotic structure and the functioning of biotic systems requires disentangling how the structural complexity of global landscapes influences the natural assemblages they can support (*20*, *21*). With more structurally complex environments supporting greater niche variety, thereby promoting species coexistence and diversification, higher structural complexity is expected to enhance species richness (*22*). However, the greater niche variety brought about by enhanced complexity can reduce the size of available niche space. Known as the Area-Heterogeneity trade-off (*23*), by reducing realised niche space, increasing structural complexity can reduce population sizes and increase the likelihood of local extirpation (*24–27*). Moreover, the term complexity is poorly defined, prompting a lack of consensus for how it is quantified and measured (*28*, *29*). For instance, *geodiversity* is a prominent measure used in quantifying abiotic environmental complexity (or heterogeneity), specifically, the richness of structural features (e.g. sediment types and landforms) and processes (e.g. riverbed erosion) comprising different environments (*15*, *30*). By synthesising the chemical, geological, and physical features comprising natural environments, geodiversity offers a proxy measure of habitat diversity that has been associated with the maintenance of ecosystem function and functional diversity (*20*, *31*). However, for identifying mechanistic drivers, there is a tension between synthesising the characteristics of underlying abiotic variables and evaluating their individual attributes.

Here, we present a quantitative global-scale assessment of structural complexity patterns using a geometric framework originally developed to quantify the structural complexity of coral reef (*32*) and forest ecosystems (*33*, *34*). Complementing efforts to classify global ecosystem types (*5*) and map the distribution of land use practices (*35*) and human occupancy (*36*), we evaluate how ecosystem and land cover classifications correspond with landscape-scale patterns in the structural complexity attributes of *height range* (*ΔH*), *rugosity* (*R*), and *fractal dimension* (*D*). Describing the shape of surfaces, *ΔH* captures the variation between maximum and minimum elevation, *R* represents a measure of true surface area, and *D* details the shape-filling capacity of a surface (*i.e.*, increases in *D* correspond with an increase in surface-contortion) (*37*, *38*). With evidence that structural abiotic environments refine local climate exposures, thus driving the establishment of similar biotic communities (*39*, *40*), we evaluate the correlation of structural attributes across differing ecosystem types. Concurrently, with anthropogenic pressures attributed to simplifying natural landscapes (*41–43*), we evaluate the relative distributions of natural environments and human modified land use types across global gradients in landscape structural complexity. Overall, we demonstrate the value of geometric descriptors of structural complexity as a unified cross-scale approach for understanding how abiotic landscape structure influences the global configuration of natural ecosystems, anthropogenic land use, and socio-ecosystems.

## RESULTS

### Quantifying global complexity patterns

Using a global terrain model sourced from the General Bathymetric Chart of the Oceans (GEBCO) (*44*) we derived estimates of height range (*ΔH*), rugosity (*R*), and fractal dimension (*D*) for the Earth’s surface at a resolution of 1.87 km, reflecting ∼185 million pixel grids. Across the globe, we report that estimates of *ΔH* and *R* align with patterns of topographic relief (Fig. 1A & B). Higher *ΔH* and *R* estimates have been shown to reflect rougher benthic surfaces in both rocky intertidal and coral reef environments (*32*, *45*). At the landscape scale, this relationship is maintained, with higher *ΔH* and *R* estimates corresponding with regions characterised by frequent tectonic activity such as the Himalayan, Andean, and Mid-Atlantic mountain ranges, and the Pacific Ring of Fire (a high density of plate boundaries encircling the coastal Pacific associated with increased volcanic and tectonic activity (*46*)). However, unlike *ΔH*, *R* also provides a representation of the space made available by increasing roughness and offers insight into finer-scale variation across surface gradients (*1*, *32*). Focusing on selected locations within the Amazon, the Mariana Trench, and the Sahara Desert (Fig. 1) illustrates how it is possible to distinguish the topographically complex canyon at the entrance to the Mariana Trench and the topographic channels associated with the Amazon River and Saharan sand dunes using just their associated rugosity values.

**Figure 1.**
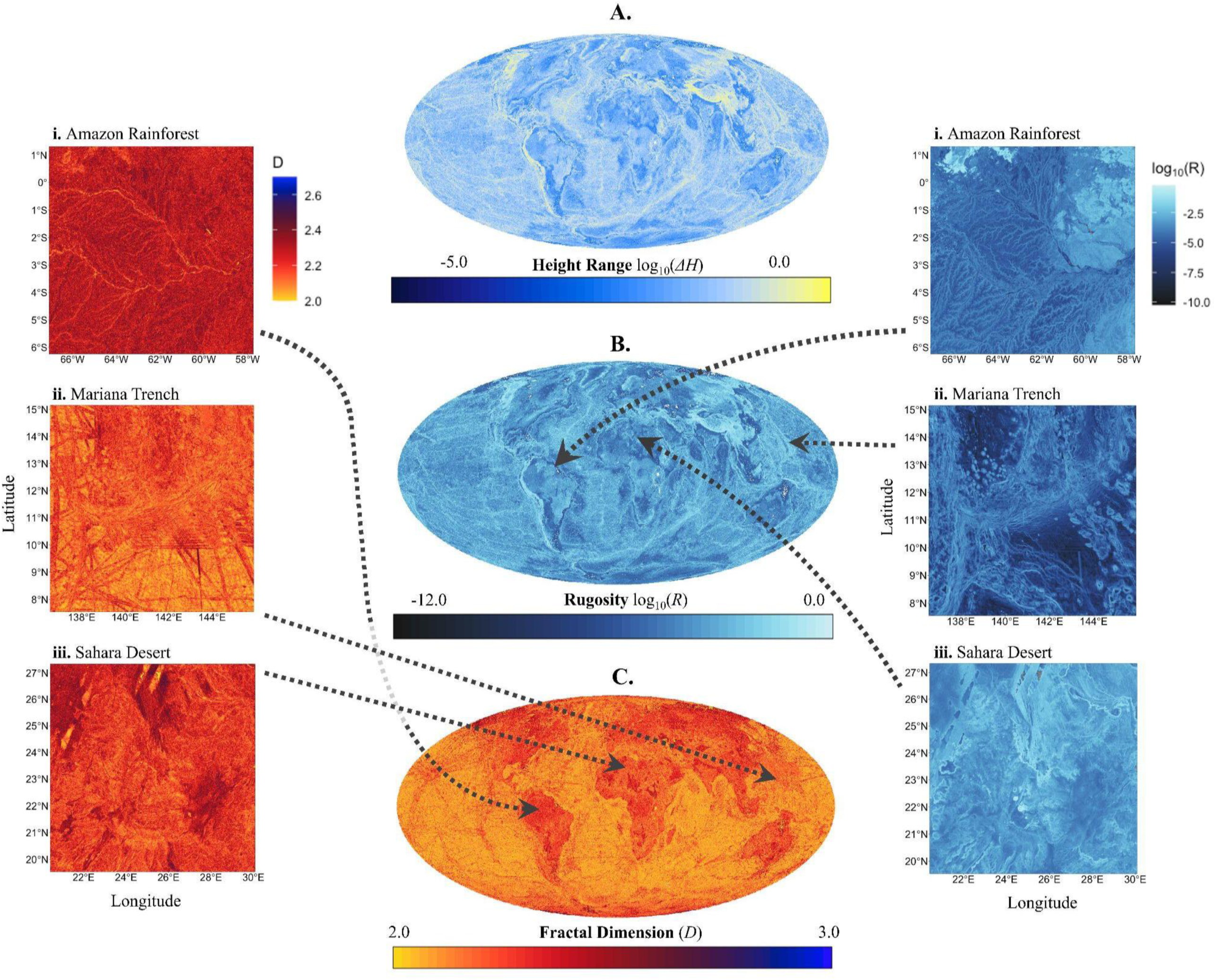
Global patterns in geometric structural complexity. High resolution maps (1.87 × 1.87 km) depicting variation in the structural complexity measures of **(A)** Height Range (*ΔH*), **(B)** Rugosity (*R*), and **(C)** Fractal Dimension (*D*), across all terrestrial and marine environments. Global maps are displayed in a global Mollweide equal area projection and comprise ∼185 million estimates of *ΔH*, *R*, and *D*, with white pixels reflecting missing or omitted entries. Comparing measures of fractal dimension and rugosity can offer differing insights into the complexity characteristics of natural environments, with the close-up panels highlighting the fractal dimension (*left*) and rugosity (*right*) of the (i) Amazonian Basin, (ii) Mariana Trench, and (iii) Sahara Desert. Each panel shows a representative 935 × 935 km section of the selected locations and is displayed in a WGS84 projection.

Areas of high fractal dimension (*D*) occur in the arctic tundras of North America and Siberia, the Amazonian and Patagonian regions of South America, and in central Africa (Fig 1C). A notable difference between the fractal properties of marine and terrestrial environments is also evident, with the global seabed typically having lower fractal dimension than the terrestrial landscape (Fig. 1C; see Appendix S2 for further details). Fractal dimension relates to the intricacy of a surface, relative to its elevation, with higher *D* suggesting increased convolution (*32*). Accordingly, differences between the fractal dimension of terrestrial and marine landscapes are consistent with the younger geological age of the marine lithosphere relative to its terrestrial counterpart (*47*) and the prominence of sedimentation throughout deep sea habitats (*48*); with, in areas of low tectonic activity, high sediment influx burying bathymetric lows and the more intricate features of marine landscapes (*49*). Geometric constraints ensure that the intricacy of a surface comes at the expense of its topographic complexity; such that increases in *D* correspond with declines in *ΔH* and *R* (when either *R* or *ΔH* are held constant) (*32*) (Appendix S1). Characterised by regions of geologically younger and less rugged terrain, rainforest river basins such as the Amazon possess high fractal dimension (Fig. 1). Alternatively, when viewed through the lens of its fractal dimension, the main canyon of the Mariana Trench almost disappears, as its surface complexity is small relative to its associated large elevational gradients (Fig. 1). Intriguingly, with higher fractal dimension often associated with an increase in habitat diversity and resource partitioning (*22*, *33*, *50*, *51*), it is perhaps not coincidental that highly diverse rainforest ecosystems are affiliated with geomorphological regions characterised by higher fractal dimension, allowing for greater habitat variety.

### Complexity and biodiversity: An island biodiversity case study

To illustrate how geometric measures of structural complexity can help distil the intrinsic association between biodiversity and abiotic structural complexity, we assessed the relationship between our complexity variables of height range (*ΔH*), rugosity (*R*), and fractal dimension (*D*), and measures of native (*i.e.*, non-introduced) island bird species and functional richness. Consistent with established theory, island area has an overwhelmingly positive effect on both island bird species and functional richness (Fig. 2A). Underlying this impact of island area, however, we evidence how the effects of structural complexity on species and functional richness patterns help to mediate the multifaceted and heterogeneous relationship between species diversity and area (Fig. 2B). Specifically, we reveal how, while increases in island rugosity and height range have little effect on bird species richness, they have a positive effect on functional diversity (Fig. 2B). The geographical barriers associated with large elevational gradients (*i.e.*, high *R* and/or *ΔH*), such as those attributed to mountainous regions, support high habitat and climatic heterogeneity, and thus are expected to promote species richness through higher rates of speciation (*6*, *19*, *52*). However, areas of high elevational complexity also support high endemism due to the specialisation required for inhabiting their associated ecosystems (*53*, *54*). On remote island systems, reductions in available niche space coupled with the specialisation needed to access and colonise island systems and tolerate the climate regimes associated with large elevational gradients, may offset any positive effect of rugosity and height range on species richness (*55*). Hence, we observe no effect of rugosity and height range on species richness but a positive effect on functional richness due to the potentially greater functional specialisation needed to inhabit these environments.

**Figure 2.**
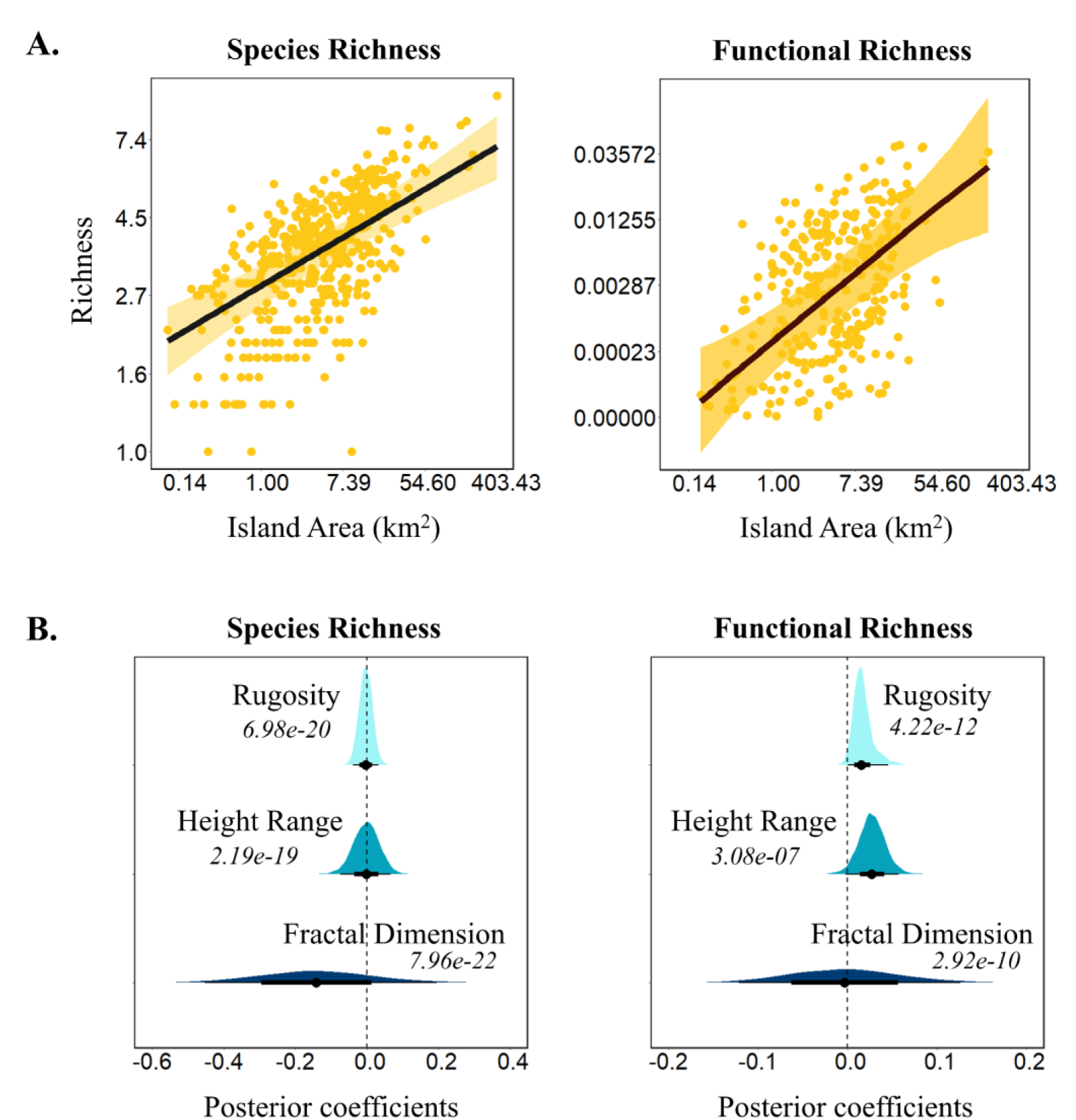
Island landscape structural complexity influences how changes in available space influence biotic configurations. **(A)** The conditional effect of Island area (km^2^) on island bird species and functional richness (*r^2^*. Species richness *=* 0.844 [95% CI: 0.828, 0.857]; Functional richness = 0.777 [0.750, 0.799]). **(B)** Posterior slope coefficients extracted from models separately evaluating the effects of the interaction between island area and island rugosity, height range, and fractal dimension, on island bird species and functional richness. Inset values show the Bayes Factor estimated for each model quantifying the effect size of including each structural complexity variable relative to the baseline scenario of only considering island area. All models account for the nested effects of archipelago and the fixed effect of island area, island latitude, distance to nearest mainland, and human population density. Error displayed as 95% CI across all panels.

Alternatively, increases in fractal dimension have a negative effect on bird species richness but little effect on bird functional richness (Fig. 2B). Higher fractal dimension across natural landscapes is thought to promote speciation through diversification by increasing habitat diversity and enabling more effective resource partitioning (*50*, *56*, *57*). However, the increased habitat diversity associated with higher fractal dimension likely reduces realised niche space, eliciting an increase in local extirpation rates (*23*). The smaller landmasses associated with remote island systems likely exacerbate any impacts of this area-heterogeneity trade-off effect on species viability. Accordingly, if island communities are close to their carry capacities, any negative effects of increasing fractal dimension would overwhelm any benefits of increasing niche variety. We note that the effect sizes associated with our observed impacts of structural complexity on bird diversity are small (Bayes Factors << 0.001; Fig. 2). The landscape scale at which we have computed our estimates of structural complexity (1.87 km) impacts our ability to isolate ecologically meaningful patterns. However, our assessment here focuses on avian fauna whose distributions are typically defined more by coarse-scale landscape features rather than fine-scale topography (*58*). Accordingly, although small, the patterns we highlight here testament the value for continued exploration into the relationship between geometric structural complexity and species diversity at finer scales across taxa.

### Structural complexity across ecosystems and land cover types

The geometric attributes associated with differing environments positions global ecosystem types along a spectrum of structural complexity, evidencing how similar ecosystem types share similar structural signatures (Fig. 3). To delineate the structural signatures associated with differing global ecosystems we isolated the distributions of *R* and *D* estimates associated with differing IUCN ecosystem functional groups (EFGs; *sensu* (*5*)). Although rugosity and fractal dimension estimates vary considerably across global EFGs (Figs. 3A, S4 & S5), these *R* and *D* distributions differ significantly among ecosystem types, with variance in structural signatures explaining a significant amount of the variance between ecosystem types (Likelihood ratio test: χ²_(df = 93)_ = 2753.04, *p* = < 0.00001). Notably, however, sequentially dropping structural variables from our models assessing these patterns suggests that independently considering the *R* and *D* signatures of different ecosystems is more informative for delimiting between ecosystem types than their additive effect; with fractal dimension the most informative of the two variables (AICs: *R + D* = 2146697, *R* = 2118558, *D* = 2001871). Marine ecosystems stand out as being characterised by a higher prevalence of less fractal and less rugose environments (Fig. 3A). Meanwhile, while transitional coastal ecosystems possess the lower rugosities of their fully marine counterparts (*59*), they exhibit higher fractal dimension (Fig 3A). Across the terrestrial realm, polar and alpine landscapes and forest and woodland ecosystems are typically more rugose environments, whilst desert and grassland ecosystems exhibit lower rugosities. Wetland ecosystems also comprise lower elevational gradients but possess very high fractal dimension. Our spectrum of ecosystem complexities also illustrates how, irrespective of their associated realm, anthropogenically modified ecosystems are typically associated with lower rugosities compared to corresponding natural environments (Fig. 3A). We note that geomorphological attributes are partly considered within the IUCNs categorisation of EFGs, raising concerns of circularity in the patterns we present here. However, this consideration only extends to the indirect mechanisms through which landscapes mediate resource availability (*60*). Focused solely on the abiotic structural attributes of ecosystem types, the patterns we illustrate here suggest a much more explicit contribution of landscape structure in configuring global biodiversity.

**Figure 3.**
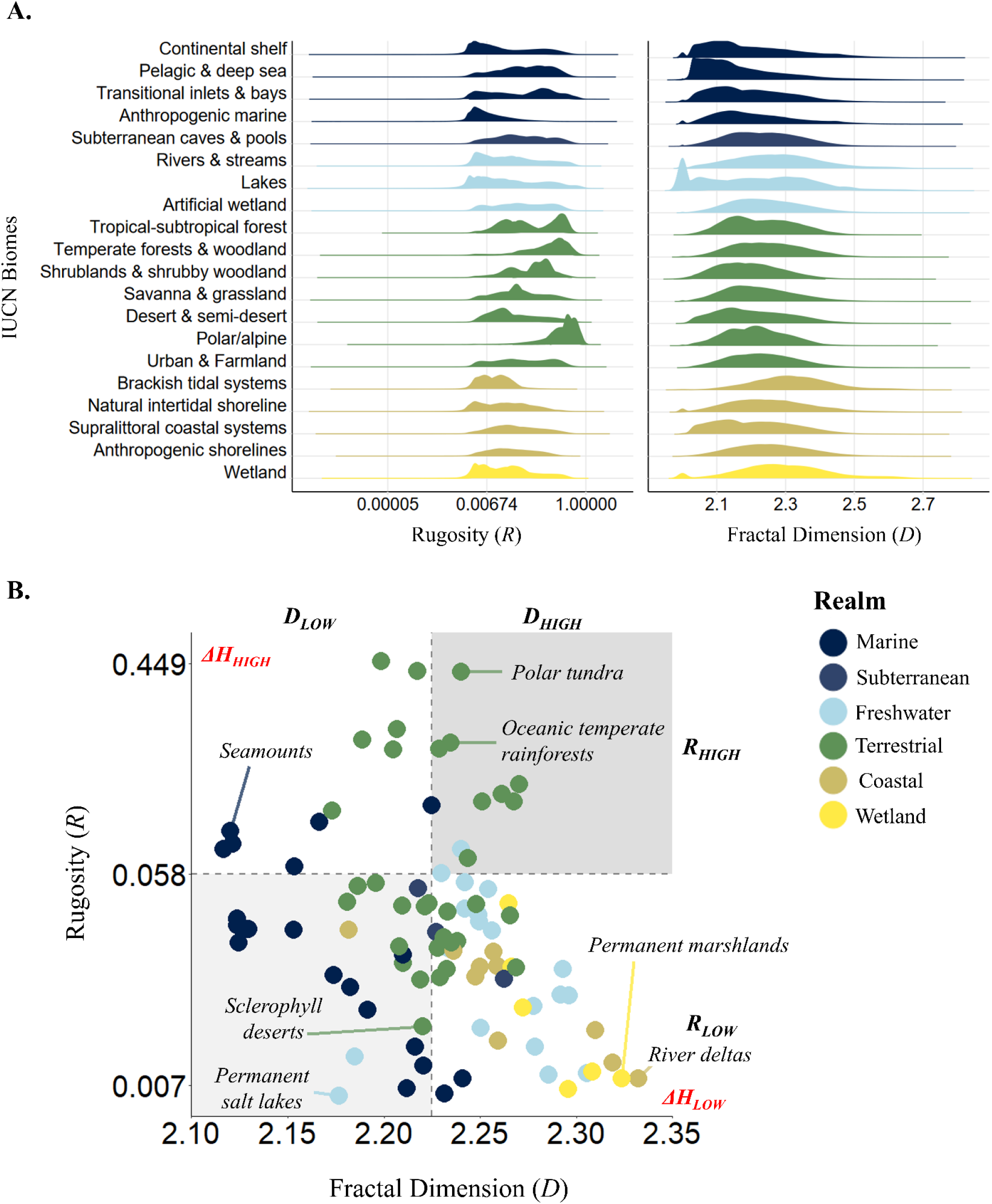
The relative structural characteristics of differing environments delineates the differing complexity regimes underlying ecosystems worldwide. (A) Ridgeline plots illustrating the distribution of rugosity (*R*) and fractal dimension (*D*) values associated with ecosystem types across different IUCN biome classifications (see Figs. S4 & S5 for a within biome breakdown of rugosities and fractal dimension across ecosystem types). Global distributions of different biomes and ecosystem types obtained from the IUCN function-based ecosystem typology (*5*). Fill colour used to delineate the realm types associated with each biome class. (B) Mean fractal dimension and rugosity estimates derived for differing ecosystem types. Point colouring used to cluster ecosystem types based on their associated realms, with panel colouring used to highlight regions of comparatively high and low rugosity and/or fractal dimension. Red inset text highlights the corresponding pattern in height range (Δ*H* across this parameter space, with representative ecosystem types shown for reference.

Correlational patterns across ecosystem complexities also offer insight into the processes involved in generating and maintaining different ecosystem types. For instance, coastal and freshwater ecosystems typically possess comparatively high fractal dimension (Fig. 3B). One exception to this pattern are permanent salt lakes (e.g. the Atacama Salt Flat), which have lower structural complexities compared to freshwater ecosystems, and instead possess similar geometric attributes to sclerophyll desert ecosystems (Fig 3B). Both permanent salt lakes and sclerophyll deserts are found in semi-arid plains (*61*, *62*), the key difference between them being the proximity of salt lake catchments to sources of periodic water inflow (*63*). Similarities in structural complexity are also apparent between permanent marshlands and river deltas (Fig. 3B). Aside from the association of river deltas with saline conditions, these ecosystem types are largely indistinguishable and possess the shared characteristics of a high deposition of silt and clay-rich sediments, constant or periodic inundation, and high turbidity (*64*). Indeed, permanent marshlands and river deltas are often components of the same environments (e.g. the Inner Niger Delta). We also highlight how oceanic temperate rainforests and polar tundra ecosystems display similar attributes of rugosity and fractal dimension (Fig 3B). Presently, these ecosystems support very different biological communities aligning with their differing climate exposures. Yet, 92 to 83 million years ago (the Turonian–Santonian age), when a far warmer global climate maintained polar temperatures > 14°C (*65*), the structural regimes associated with contemporary polar tundra supported diverse temperate rainforests (*66*).

Further exploring how the structure of natural landscapes influences the biotic assemblages they support, we compared the different but complementary perspectives of geometric structural complexity and geodiversity, which represents the combined richness of geological, geomorphological, pedological, and hydrological features comprising different environments. Aligning global maps of terrestrial geodiversity with our global maps of structural complexity, we observed how environments comprising a greater richness of different geological and hydrological features typically exhibit higher rugosities and greater elevational gradients (Fig. 4A & B). Alternatively, fractal dimension and geodiversity exhibit no apparent relationship (Fig. 4C). Fundamentally, fractal dimension and geodiversity reflect different continuous and categorical measures of geomorphological variability, and so it is to be expected that they would portray different patterns. However, the same is also true of geodiversity and both rugosity and height range. With geodiversity now a well-established measure used in quantifying how patterns in environmental heterogeneity influence biodiversity, we illustrate how geodiversity and fractal dimension reflect orthogonal descriptors providing differing insights into the same abiotic landscapes. Although increases in both geodiversity and fractal dimension have been reported to increase biodiversity (*22*, *67–69*), often their effects on diversity are complex and non-linear (*70*). Consequently, it would be worthwhile to explore how the interaction between a landscape’s abiotic richness and its geometric complexity impacts resource availability and how organisms exploit them.

**Figure 4.**
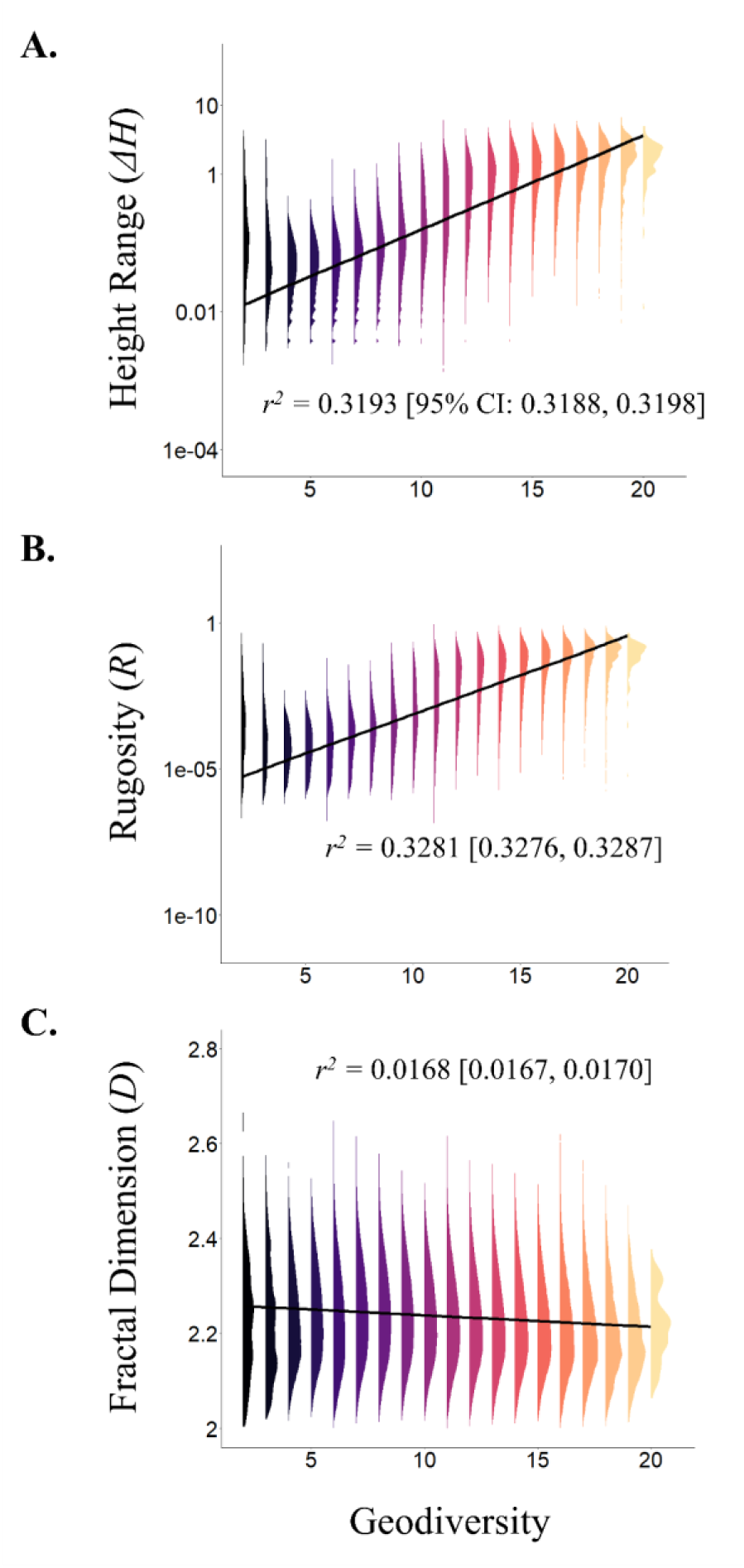
Comparing the geometry and structural richness of natural environments allows for disentangling the mechanisms through which structural complexity mediates the configuration of global biodiversity. The alignment of the structural complexity measures of (A) Height Range (*ΔH*), (B) Rugosity (*R*), and (C) Fractal Dimension (*D*) with estimates of geodiversity (geology, geomorphology, pedology, and hydrology combined). Geodiversity estimates were extracted from a high resolution (1.87 × 1.87 km) global map depicting patterns in the richness of structural features comprising an environment (*117*). The solid line across each plot represents the predicted relationship between geodiversity and each complexity measure.

Finally, our evaluation of global structural complexity patterns suggests a nuanced relationship between anthropogenic activity/human occupancy and the complexity of natural landscapes (Fig. 5). We quantified the rugosity and fractal dimension patterns associated with the global distributions of different land cover classifications obtained from the Copernicus Climate Change Service (C3S) land cover survey (*35*). Rugosity and fractal dimension estimates vary considerably across global land use types (Fig. 5A). Again, however, variance across the *R* and *D* signatures associated with different land use types explains a significant amount of the variance among land use categories (Likelihood ratio test: χ²_(df = 13)_ = 1160.54, *p* = < 0.00001). Stepwise variable omission then indicates that the interaction between *R* and *D* is more important for distinguishing between land use categories than either variable independently (AICs: *R + D* = 189848, *R* = 190982, *D* = 196095). For instance, although clustering rugosity and fractal dimension estimates by associated land cover categories represents a coarse filter, it is still apparent how human modified habitats (*i.e.*, urban and cropland environments) typically display lower complexities compared to more natural land cover types, apart from naturally bare substrate and permanent snow and ice (Fig. 5A). Our findings also suggest that while human occupancy likelihood is largely unaffected by changes in structural complexity, human population densities appear more associated with the structural complexity of natural environments (Fig. 5B-D). Although structural complexity alone is a weak predictor of human population densities (*r^2^* < 0.011). Overlaying gridded human population densities sourced from NASA (*36*) onto our maps of global complexity we evaluated how human population densities worldwide correspond with patterns in *ΔH*, *R*, and *D*. Specifically, while rugosity and height range are positively correlated with human population densities, increases in fractal dimension correspond with large declines in human population density (Fig. 5D). Note, human population density was power transformed due to extreme skew, so a change in density values from 1.08 to 0.95 reflects an order of magnitude shift in people/km^2^.

**Figure 5.**
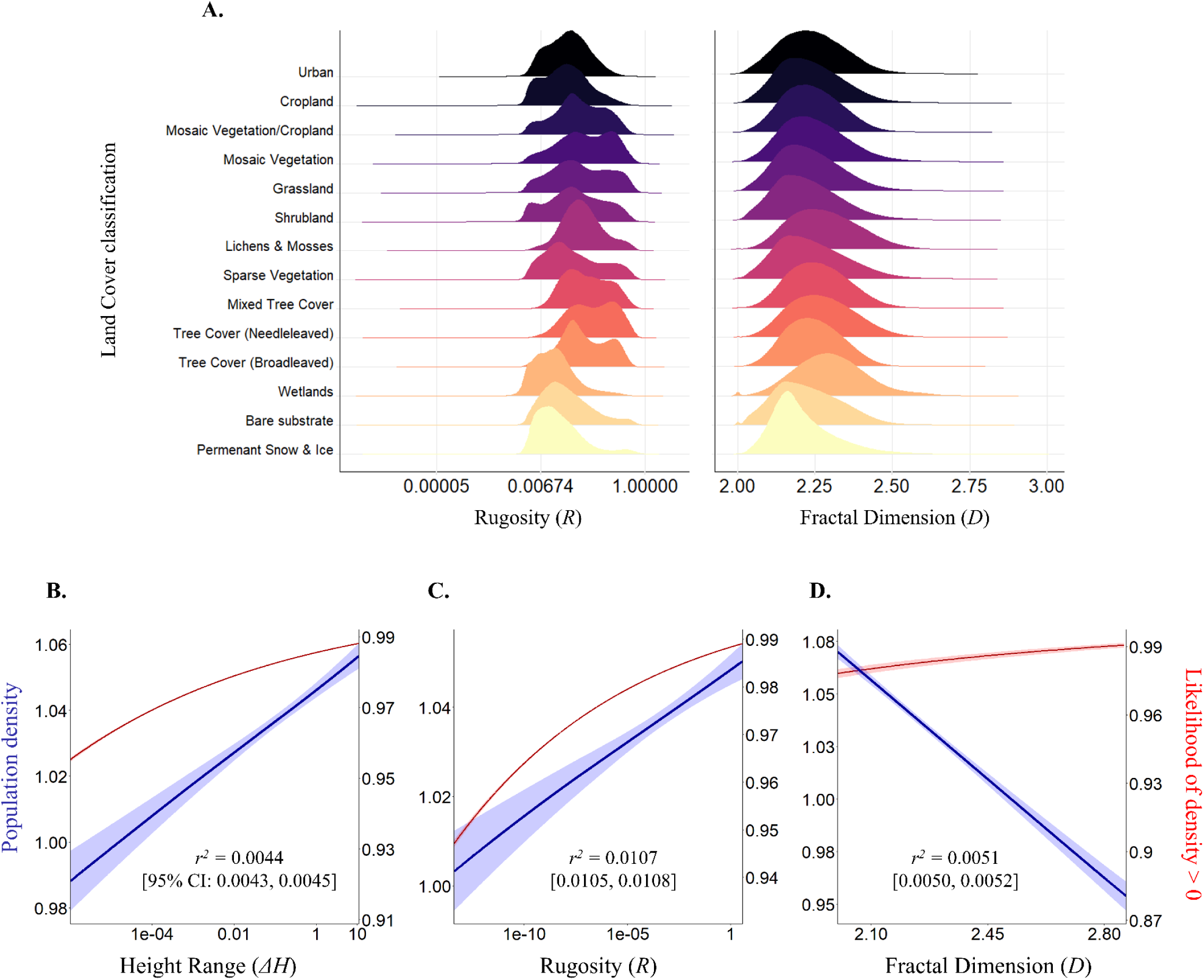
Global patterns in structural complexity influence how humans interact with and use natural environments worldwide. **(A)** Ridgeline plots illustrating the distribution of rugosity (*R*) and fractal dimension (*D*) estimates associated with different land cover classifications. Land cover categories and their spatial distributions obtained from the Copernicus Climate Change Service (C3S) land cover survey (*35*). **(B-D)** The alignment of the structural complexity measures of **(B)** Height Range (*ΔH*), **(C)** Rugosity (*R*), and **(D)** Fractal Dimension (*D*) with patterns in human population density. Human population density patterns were sourced from the Gridded Population of the World (*36*), with each plot showing the predicted relationships (including 95% CI) derived from two-part hurdle models separately exploring patterns in the probability of greater than zero population densities (red) and recorded population count densities (blue). Population density estimates were transformed using a power transformation (*y^0.05^*).

## DISCUSSION

Geophysical processes are key to understanding the global and regional distribution of biodiversity (*19*). We demonstrate here how using geometric measures to quantify cross-system patterns in global structural complexity can inform our understanding of the processes through which structural complexity affects abiotic and biotic regimes. Understanding the drivers of biodiversity, and how natural communities respond to ongoing global change, requires an appreciation for the intrinsic association between biodiversity and abiotic structural complexity (*71*). Colonisation, extinction, and speciation are fundamental drivers of global patterns in species richness (*72*, *73*) and are influenced by the structural architecture of natural landscapes (*7*). Moreover, landscape morphology and structure mediate whether, how, and when, organisms can access available resources (*6*). Indeed, by shaping dispersal pathways and niche partitioning trends, historic shifts in landscape structure are thought to have influenced the contemporary arrangement of global biodiversity more so than prevailing climatic regimes (*52*, *71*, *74*, *75*).

Habitat theory argues that whilst climatic factors are the key determinants of speciation and diversification, neither speciation nor diversification can happen without some extrinsic forcing from the physical environment (*7*). Our approach does not allow for distinguishing whether structural complexities determine ecosystem types, ecosystem forming processes determine structural signatures, or whether both are co-determined by climatic or tectonic history. However, conveying associations between landscape structural complexity and differing ecosystem types illustrates the value of using geometric approaches as a complementary approach to disentangle the mechanisms delineating natural ecosystems and biodiversity. The Amazonian Basin, considered amongst the planet’s most biodiverse ecosystems (*76*, *77*), exists in a terrain with a high fractal dimension, reflecting its riverine geometry. With topographic barriers capable of both inducing local extinction and creating regional refugia, climate shifts across more complex landscapes will impose more frequent selection pressures and fragmentation (*7*, *19*); resulting in greater species richness. Likewise, increases in fractal dimension also support greater species coexistence through increased resource partitioning (*50*, *56*, *57*). Nevertheless, the Saharan Desert landscape, a comparatively more inhospitable region, also possesses high fractal dimension. Yet rather than weaken the argument that structural complexity plays a role in configuring biodiversity, the biotic components of natural ecosystems are typically more transient through time than topographical features (*2*). Although presently species poor relative to the Amazon rainforest, under differing past climate regimes, the Sahara Desert supported vibrant grassland communities (*78*); evidencing a potential mechanism through which structural complexity can mediate how local climate regimes configure global biodiversity patterns through time.

Alternatively, high fractal dimension across seemingly unproductive landscapes is hypothesised to allow associated systems to support higher than expected levels of biodiversity (*80*). Coexisting species cannot partition available space indefinitely (*81*). However, cross-scale patterns in fractal dimension (*i.e*., the shape of valleys and ridges down to the interstitial space between sand particles) allow different species from similar trophic niches to harvest the same resources within the same habitats (*82*). Although geometric complexity patterns observed in coral reef and forest ecosystems (*32–34*) persist at the landscape-scale, how organisms respond to these patterns is scale-dependent (*83*). The size and mobility of organisms determine how they use their environments and thus their perception of any associated complexity and fragmentation (*22*, *83*). Indeed, the negative effects of declines in realised niche space associated with increasing fractal dimension are expected to be amplified at finer spatial scales (*22*). With scale a major factor in determining the effect of observed patterns in structural complexity and how they are interpreted, establishing a definitive understanding of the ecological role of abiotic structural complexity requires a multiscale perspective (*38*, *83*). The global scope of our approach here necessitated working at a coarse landscape scale resolution (1.87 km). However, with ecologically relevant patterns still manifesting at this scale, we illustrate the value of geometric complexity measures for establishing cross-scale understanding of the processes helping to shape biodiversity patterns worldwide.

The relationship between species diversity and area has long been central to our understanding of biodiversity theory (*84*), yet environmental context underpins how the relationship actually manifests (*85*, *86*). Applying our quantification of structural complexity to the assessment of avian species-area relationships observed across global island systems highlights the nuanced insights structural measures can offer into the mechanisms mediating the configuration of global species and functional diversity patterns. Area is important for providing the space necessary for species to coexist (*73*, *87*, *88*). Whether changes in available area constitute increases or decreases in rugosity, height range, or fractal dimension will refine how these changes actually influence the configuration of local species diversity patterns. Using geometric measures of structural complexity can allow for quantifying mechanisms through which changes in area influence biotic communities (e.g. changes in habitat diversity, niche size and/or species isolation). Accordingly, using these measures we can reconcile understanding of the nuanced effects of area on species diversity patterns across ecosystems and taxa.

Beyond its effects on biodiversity, the complexity of natural landscapes profoundly impacts multiple aspects of human society (e.g. tourism, agriculture, mobility, and wellbeing) (*2*, *18*). For instance, intuitively, with mountain ranges and high alpine regions typically associated with a reduction in the space and arable land needed for agriculture and food production, the development of human settlements has often gravitated towards flatter, more lowland environments (*89*). Accordingly, we observed how environments exposed to the greatest levels of anthropogenic pressure and exploitation typically possess reduced topographical complexity. Yet, we also report how structural complexity has little effect on human occupancy patterns. Any expectation that humans select less complex environments for development oversimplifies the technological and socioeconomic drivers of anthropogenic land use (*18*, *90*). Instead, the contemporary relationship between human society and landscape complexity is likely more impacted by our capacity for modifying landscapes to create less complex environments, with, for example, agriculture flattening natural terrains and urbanisation homogenising landscapes (*41–43*, *91*, *92*). While we are unable to distinguish between cause and effect with our static layers, the two processes are not mutually exclusive, and both instances are likely true across the global landscape. Indeed, this tautological relationship is apparent in our observation that more fractal environments appear to support lower human population densities. Rather than indicating human populations to be adversely affected by reduced niche space (*i.e.,* the area heterogeneity trade-off), the satellite-derived DEM underlying our complexity calculations incorporates anthropogenically modified environments, thus many regions displaying low fractal dimension likely do so because they have been modified to accommodate large human populations.

Our global-scale assessment highlights the role that geometric structural complexity plays in mediating the configuration of natural and anthropogenic communities. Having long appreciated the association between diversity and area (*33*, *84*), a recent focus of ecological research has been to quantify the mechanistic drivers linking biodiversity and the physical structure of natural environments (*14*, *20*, *72*). Nature also does not exist in a vacuum, with the majority of global ecosystems facing some form of anthropogenic pressure (*93–95*). How we interact with and use our local environment is mediated by patterns in their structural complexities. So, too, is how these natural environments respond to our activities. The use of geometric theory to quantify and isolate patterns in the structural complexity of global environments can provide continuous quantitative variables for delineating ecotones based on their shared structural characteristics. Moreover, we have demonstrated how a geometric framework developed for reef- and forest-scale applications represents a unifying approach for quantifying structural complexity and its effects on biodiversity across scale. Overall, our approach offers insight into the future configuration of global biodiversity by serving as a basis for evaluating how structural complexity helps to shape biotic communities, and how this association will be influenced by anthropogenic pressures and ongoing global change.

## MATERIALS AND METHODS

Our study comprised the following sequence of steps: [1] generating global maps of selected complexity indices; [2] assessing the value of including structural complexity variables in tests of island diversity-area relationships; [3] evaluating ecosystem and land-use associated complexity characteristics; and [4] quantifying the association between structural complexity and selected abiotic and biotic mechanisms. All spatial analyses were carried out using R (*96*) and QGIS (*97*), with our code archived on GitHub (https://github.com/CantJ/Global-Geometric-Complexity) for download or cloning.

### Generating global complexity maps

To evaluate patterns in the structural attributes of global land- and seascapes, we applied a geometric framework used in quantifying the structural shape of planar surfaces, to a global ocean and land terrain model (including ice surface elevation) sourced from the General Bathymetric Chart of the Oceans (GEBCO; https://www.gebco.net/) (*44*). Specifically, we focused on the geometric measures of height range (*ΔH*), rugosity (*R*), and fractal dimension (*D*). Initially we merged 15 arc-second resolution terrain raster tiles (EPSG: 4326, WGS84) downloaded from GEBCO to produce a global map of surface elevation in metres. To ensure a consistent resolution across raster cells, we reprojected this elevation model (henceforth digital elevation model [DEM]) to an equal-area Mollweide projection (EPSG: 54009) using bilinear resampling. Note, the native reprojection (*i.e.*, the use of default parameters) of the WGS84 GEBCO DEM to an equal-area Mollweide projection results in a resolution of 186.754 m. Thus, to simplify our subsequent computation of geometric indices, during this reprojection we imposed a minimum resolution of 187 m (*L0*). From this reprojected DEM, using the associated functions contained in the *habtools* package (*98*) we then extracted estimates of *ΔH*, *R*, and *D*, at a scale of 1870 m (*L*; one order of magnitude greater than *L0*, see Appendix S3 for scale sensitivity analysis). Briefly, this extraction involved separating our DEM file into a series of *L × L* grids, with each grid therefore comprising 100 raster cells (*i*.*e*., 10 *×* 10 cells). Using multicore processing, implemented using the *foreach* (*99*) and *doParallel* packages (*100*), we iteratively processed each grid to extract estimates of *ΔH*, calculated as the difference between the highest and lowest elevation in each grid, and *R*, defined as the surface area of each grid over its planar area. We computed *R* using the *habtools* ’area method’, following the approach outlined in Jenness (*101*) which entails generating 3-dimensional triangles linking the centroid of each grid cell to the centroids of all adjacent cells before estimating the area of each triangle laying within the focal grid cell boundary. After computing ΔH and R, we then calculated *D*, for each grid, by inputting corresponding *ΔH* and *R* values into the geometric plane equation outlined in Torres-Pulliza *et al.* (*32*) (equation 1). Hence, we computed *D* using an approach referred to as the height variation method.

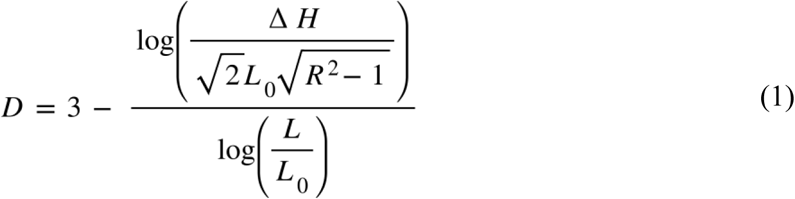

Next, we carried out data cleaning, omitting all estimates that violated geometric theory for surface habitats (*ΔH* < 0; *R* < 1; 2 > *D* > 3; 85,183 entries, 0.046% of initial sample) and/or corresponded with non-fractal (flat) grids (*ΔH* = 0, *R* = 1 & *D* = 2; 435,225 entries, 0.233% of initial sample). We omitted non-fractal grids on the basis that, at a resolution of 1870m, perfectly flat landscapes do not exist and instead represent artefacts present in the original elevation model (*49*), or accuracy errors introduced during model reprojection. Prior to collating our estimates for each complexity measure into separate spatial raster files we transformed all *ΔH* and *R* estimates according to their expression in the geometric plane equation (*32*), such that rugosity is expressed as *R*^2^ – 1 and height range is expressed as .

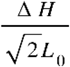

### Disentangling drivers of island biodiversity

#### Quantifying avian diversity

To assess the role structural complexity plays in helping to mediate the configuration of biological communities, we modelled the independent relationships between the complexity measures of *ΔH*, *R*, and *D*, and bird species diversity, within the context of assessing island species-area relationships (ISARs). We sourced records of observed species lists (presence data) from island bird surveys carried out on 476 islands across 49 island archipelagos (*102*, *103*) (Table S3). These species lists focus on native (*i.e.*, non-introduced) land bird species, thus seabird species were excluded, and all species counts pool together endemic and cosmopolitan species. Using these species lists we estimated counts of avian species richness for each island as the total count of observed species (n = 476 counts). Next, for each species recorded across the island species lists, we complied details describing 11 traits related to beak size (tip to skull length, tip to nares length, width, and depth), wing size (wing length and hand-wind index), body size (tarsus length, tail length, and body mass), and foraging ecology (trophic niche and locomotory foraging niche) from the AVONET database (*104*). With these trait values, we then used the *mFD* package (*105*) to compute the functional distances between each species (calculated as Gower distances due to the inclusion of non-continuous traits), and subsequently the functional richness of the bird communities recorded on each island (n = 311 counts). We then applied transformations to species richness (*log10*) and functional richness (*y^0.275^*) prior to further analysis.

#### Quantifying island complexity

The Global Island Database maintains a series of shapefiles detailing the location and size of > 340,000 islands worldwide (*106*). From this resource we extracted the planar area (*AI*; km^2^), the distance to the nearest mainland (Euclidean distance, metres), and the centroid latitude, for each island for which we had bird species and functional richness estimates. We also used the corresponding island shapefiles to isolate the structural complexity signatures (*ΔH*, *R*, and *D*) associated with each island from the aforementioned global maps. Likewise, we also extracted the associated human population densities (*PD*) for each island from a map of the global population density recorded in 2020 sourced from the Gridded Population of the World (*36*) and reprojected to a 1870m resolution Mollweide projection. Using these isolated values, we were then able to calculate mean complexity values and total *PD* for each island. We applied a *log10* transformation to our variables of island area and *PD* (*PD*+1), and a power transformation to our island isolation estimates (*y^0.5^*). Finally, we converted centroid island latitude estimates into measures of absolute latitude.

#### Modelling species-area relationships

Independently, for our measures of avian species richness and functional richness (henceforth ‘richness’), we implemented a series of Bayesian regression models (equation 2) separately modelling the relationships between our estimates of richness (*y*) and island area (km^2^), island rugosity, island height range, and island fractal dimension (each represented by *x*):

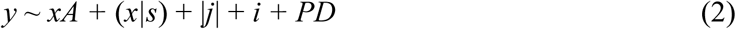

To account for spatial autocorrelation across island chains we included the nested effects of island archipelago (*s*). Each model also included the fixed covariates of absolute latitude (*j*), distance to nearest mainland (*i*), and *PD*, to account for the confounding effects of the latitudinal biodiversity gradient, the impact of island isolation on colonisation, and exposure to anthropogenic pressures. To account for the interaction between area and structural complexity, we also included island area (*A*) as a fixed effect in our models exploring the relationship between richness and island rugosity, height range, and fractal dimension. Apart from our model quantifying the relationship between functional bird richness and island area (*FR ∼ A*), preliminary ‘leave-one-out’ (LOO) model cross-validation showed no significant difference between the expected log predictive densities (ELPD) associated with applying a linear or polynomial modelling approach (ELPD. *FR ∼ A*: polynomial >> linear; polynomial = linear in all other cases). From each corresponding model, we extracted the standardised posterior slope coefficients quantifying the relative effects of rugosity, height range, and fractal dimension on island bird species and functional richness. Across our sets of species and functional richness models, we also computed Bayes Factors quantifying the effect size of each structural complexity variable on bird richness relative to the baseline of only considering island area. All modelling for this island biodiversity case study was carried out using the *brms* package (*107*), with each model initiated using uniformed priors and implemented over four Markov chains each consisting of 3000 iterations.

### Comparing ecosystem and land cover complexities

We evaluated patterns across our estimates of *R* and *D* associated with differing ecosystem and land cover classifications. Carrying out this assessment involved assigning each grid cell across our complexity maps to particular ecosystem and land-use types. Firstly, we used the IUCN function-based ecosystem typology (5) to define ecosystem types and their spatial distributions across the global landscape. Briefly, this hierarchical classification collates ecosystems into Realms, Biomes, and Ecosystem Functional Groups (EFGs), based on shared functional characteristics, rather than associated species assemblages. For instance, a representative biome within the terrestrial realm is the savannas and grasslands biome. This biome consists of five EFGs, grouped by their shared exposure to strong seasonal cycles in water availability and maintenance of highly productive, continuous ground layer grass communities. These five EFGs are then distinguished between based on: the biotic structure associated with their dominant foundation species (Hummock savannas & Temperate woodlands) (*108*, *109*), their reliance on recurring fires for maintaining coexistence (Pyric savannas) (*110*) or large herbivores for maintaining heterogeneous vegetation (Trophic savannas) (*111*), and their climate exposures (Temperate sub-humid grasslands) (*112*).

From the IUCN ecosystem typology, we sourced a series of global maps detailing the differing spatial distributions of all ecosystem classifications (*113*). Next, we quantified spatial patterns in land-use categories using satellite-derived land cover data for 2020 sourced from the Copernicus Climate Change Service (C3S) (*35*). From the C3S data portal, we sourced an .nc file comprising different global land cover variables derived from satellite surveys carried out in 2022. From this file we extracted the associated land cover class layer using the *raster* package (*114*). Finally, we reprojected all ecosystem type- and land cover-maps to a Mollweide projection (EPSG: 54009) at a resolution of 1870m (*L*), whilst ensuring finalised map extents corresponded with that of our original DEM. For the assessment of both ecosystem and land-use associated complexities we overlayed the corresponding spatial files detailing the spatial distributions of selected ecosystem and land cover types (Tables S1 & S2) onto our global complexity maps to isolate the distribution of *R* and *D* estimates associated with each ecosystem and land cover category. We then used paired multinomial logistic regression and likelihood-ratio tests to evaluate whether the structural complexity signatures associated with different ecosystem and land cover types quantitatively discriminate among ecosystem and land cover types. Isolating a randomly selected 1% sample of the *D* and *R* estimates associated with each ecosystem type and land use category (total sample size: Ecosystem = 375,460 across 96 levels, Land use = 41,445 across 14 levels), we implemented multinomial logistic regression separately quantifying the differences among ecosystem types and land use categories as a function of their differing structural complexity signatures (*y* ∼ *D* + *R*). Next, we specified null models simply quantifying the differences among ecosystem types and land use categories (*y* ∼ 1), before using a likelihood-ratio test to determine whether including the structural metrics explains more variance among ecosystem types and land use categories than these null models. We then explored the effect of sequentially dropping the *D* and *R* variables from our models, using model AICs to compare subsequent model strength with the fully specified version. We implemented our multinomial logistic regression and likelihood-ratio tests using the *nnet* (*115*) and *lmtest* (*116*) packages, respectively. Finally, to aid visual comparison of the different structural complexity signatures associated with differing ecosystem functional types, from these distributions of *R* and *D* values we also computed ecosystem-level mean estimates.

### Evaluating associated global variables

Focusing on the terrestrial landscape, we evaluated how our estimates of *ΔH*, *R*, and *D* correspond with two established drivers of global biotic and abiotic regimes: geodiversity (*G*), and human population density (*PD*). We sourced geodiversity estimates from a global map of terrestrial geodiversity developed during an assessment of geodiversity representation across UNESCO global geoparks (*117*) (EPSG: 54009). Originally, sourced at a resolution of 250m, this global map was reprojected to a resolution of 1870m (*L*) and to an extent equal to that of our global complexity raster layers. Meanwhile, we obtained global data on human population densities from the Gridded Population of the World (*36*). We downloaded a map of the global population density recorded in 2020 at a 30 arc-second resolution (EPSG: 4326) which we subsequently reprojected to a 1870m resolution in a Mollweide projection. Aligning corresponding grid values across these spatial maps, then allowed for extracting corresponding estimates for *ΔH*, *R*, *D, G*, and *PD*, from which we retained the grids with complete entries for all variables (∼38 million grids). Following data omission, we used the *terra* package (*118*) to test each of our structural complexity variables (*ΔH*, *R*, and *D*) for spatial autocorrelation, by computing Moran’s *I* statistic for each measure (*I*: *ΔH* = 0.883; *R =* 0.805; *D =* 0.476, assuming a Queen’s case nearest neighbour matrix, *i.e.*, all adjacent grid cells including diagonals). Lastly, we transformed our population density variable for normality. This transformation took the form of a power transformation (*y^0.05^*) applied only to non-zero entries.

We analysed the pairwise relationships between each of our geometric variables (*ΔH*, *R* and *D*) and *G*, using Bayesian regression models implemented with uninformed priors using Integrated Nested Laplace Approximation (INLA) and Monte-Carlo resampling. To assess each pairwise relationship we ran our models using the *qbrms* package (*119*), repeating each pairwise comparison across 100 iterations using a randomly selected sub-sample of 100,000 data points; each time computing the associated model’s *r^2^* value and predicted relationship. We fitted all models using a linear format set to a gaussian distribution, with geodiversity set as the predictor variable, and with latitude and longitude included as continuous fixed variables to account for the spatial autocorrelation evident across our complexity variables (*I* > 0). In all cases, as the less complex modelling format, linear modelling was selected over a polynomial approach with LOO model comparisons showing no significant difference between either approach (ELPD. polynomial = linear in all cases).

Also implemented using Monte-Carlo resampling, we analysed the pairwise relationships between each of our geometric variables and *PD*, using manually specified hurdle models again implemented with uninformed priors using INLA with the *qbrms* package. Using this hurdle approach allowed us to separately model the probability of zero population density as a function of structural complexity, and the relationship between non-zero population densities and structural complexity. We repeated each pairwise hurdle model 100 times on a randomly selected sub-sample of 100,000 data points, each time retaining the model’s *r^2^* value for its non-zero component, predicted non-zero relationship, and hurdle coefficients (probability of being zero). We modelled the probability of zero population density as a function of our complexity variables using a binomial distribution, and the relationship between complexity and non-zero population densities using a gaussian distribution. Preliminary LOO model comparisons indicated no significant difference in applying a linear or polynomial format to the modelling of non-zero population densities (ELPD. polynomial = linear in all cases), with a linear approach selected as a result. Again, all models included latitude and longitude as continuous fixed variables to account for spatial autocorrelation across our complexity variables.

## Author Contributions

Conceptualization: JC, NS, EMPM, JSM, MD

Methodology: JC, NS, ACS, KFR, JSM, MD

Investigation: JC

Visualization: JC, JSM, MD

Funding acquisition: JC, JSM, EMPM, MD

Project administration: MD, JSM

Supervision: MD

Writing – original draft: JC, MD

Writing – review & editing: JC, NS, EMPM, ACS, KFR, JSM, MD

## Data, code, and materials availability

All data and code needed to evaluate and reproduce the results in the paper are present in the paper and/or the Supplementary Materials. This study did not generate new materials. The raw data, code, and described global complexity maps have been archived open access on Zenodo (https://doi.org/10.5281/zenodo.15006508).

## Supplementary Materials

Supplementary Text

Figs. S1 to S5

Tables S1 to S3

References (120–121)

