## Appendix for "Landscape structural complexity influences the global distribution of ecosystems and people"

Science Advances
AAAS

### Supplementary Materials for

#### Landscape structural complexity influences the global distribution of ecosystems and people

James Cant *et al.*

**The PDF file includes:**

Supplementary Text
Figs. S1 to S5
Tables S1 to S3
References (120 - 121)

#### Supplementary Text

##### S1. Quantifying geometric relationships

Aligning corresponding grid cell values across our global complexity maps, we quantified the pairwise relationships between $\Delta H$, $R$, and $D$, using a Bayesian modelling framework combined with Monte-Carlo resampling. We carried out these models using the *qbrms* package (*119*). Across pairwise comparisons, we modelled the relationship between $\Delta H$ and $R$ using a polynomial distribution whilst we modelled the relationships between $\Delta H$ and $D$, and $R$ and $D$ using a linear distribution. Model distributions were selected based on differences in expected log predictive densities (ELPD) computed using 'leave-one-out' (LOO) cross-validation model comparisons (LOO ELPD. $\Delta H$ *vs* $R$: polynomial >> linear; $D$ *vs.* $\Delta H$: polynomial = linear; $D$ *vs* $R$: polynomial = linear). In each case, our regression framework included latitude and longitude as continuous fixed variables to accommodate for the positive spatial autocorrelation evident across each complexity variable. We repeated each pairwise comparison 100 times, each time selecting a random sample of 100,000 data points before implementing a Bayesian regression model fit with uninformed priors using integrated Nested Laplace Approximation. Assessing the resultant pairwise relationships between our estimates of $\Delta H$, $R$, and $D$ affirmed a positive correlation between $R$ and $\Delta H$ ($r^2$ = 0.97139 [95% CI: 0.97135, 0.97143]; Fig. S1A). Meanwhile, the relationship between $D$ and both $R$ and $\Delta H$ is more nuanced. As such, as a surface's fractal dimension increases, then either its height range decreases ($r^2$ = 0.08715 [0.08674, 0.08757]; Fig. S1B) or its rugosity increases ($r^2$ = 0.01767 [0.01748, 0.01787]; Fig. S1C), or both.

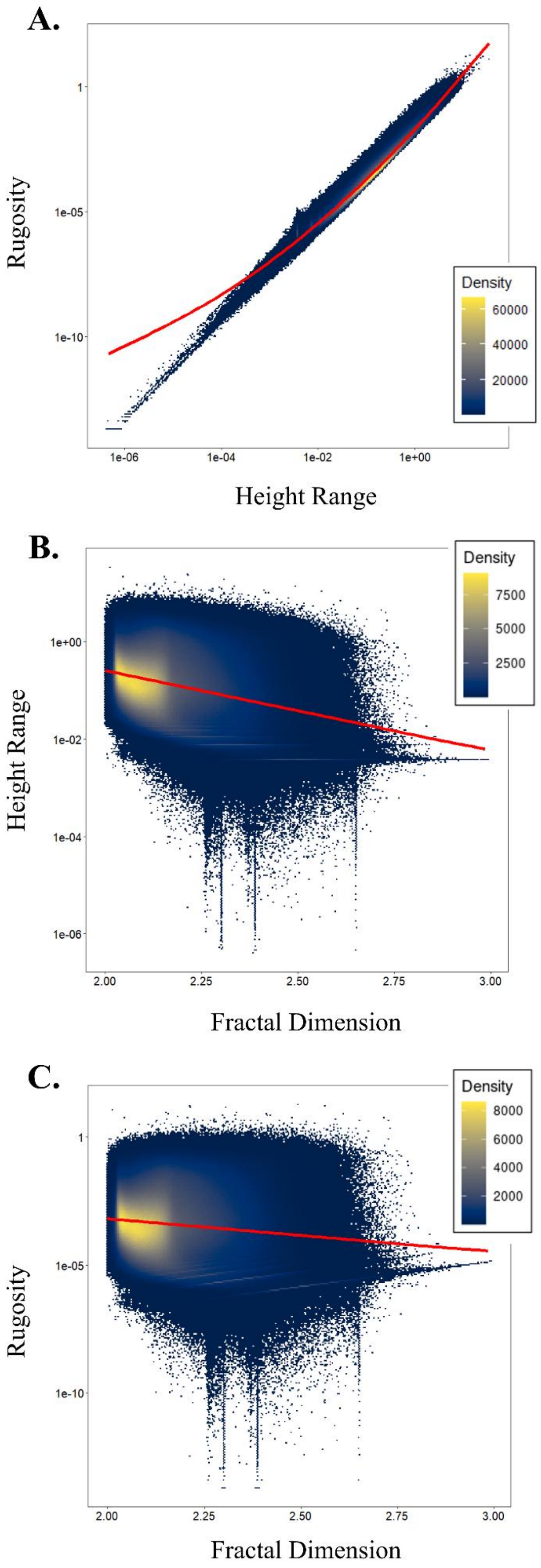
A.
Rugosity
1
1e-05
1e-10
Height Range
1e-06
1e-04
1e-02
1e+00
Density
60000
40000
20000
B.
Height Range
1e+00
1e-02
1e-04
1e-06
Fractal Dimension
2.00
2.25
2.50
2.75
3.00
Density
7500
5000
2500
C.
Rugosity
1
1e-05
1e-10
Fractal Dimension
2.00
2.25
2.50
2.75
3.00
Density
8000
6000
4000
2000

**Figure S1. Fundamental geometric constraints maintain the pairwise relationships between the complexity variables of Height Range ($\Delta H$), Rugosity (*R*), and Fractal Dimension (*D*) at the global scale.** Plots depicting the relationships between **(A)** *R* & $\Delta H$, **(B)** $\Delta H$ & *D*, and **(C)** *R* & *D*. Plotted points (n = 20 million) represent a ~14% sample of the global dataset, with point colour shades corresponding with the density of overlapping points. The solid red line drawn across each plot represents the predicted relationship between each variable pair quantified following 100,000 resampling iterations of a Bayesian regression framework including fixed spatial random effects.

S2. Comparing land- and seascape complexities

To evaluate for differences among our estimates of *ΔH*, *R*, and *D* between marine and terrestrial environments, we assigned each grid cell across our complexity maps as either terrestrial or marine. We identified whether grid cells were associated with either the terrestrial or marine realm using a terrestrial land mask generated using land cover data from the Copernicus Climate Change Service (C3S) (*35*). From the C3S data portal, we sourced an .nc file comprising different global land cover variables derived from satellite surveys conducted in 2022. From this file, we extracted the associated land cover class layer using the *raster* package (*114*), which we then reprojected to a Mollweide projection (EPSG: 54009) at a resolution of 1870m (*L*), all whilst ensuring the finalised map extent corresponded with that of our DEM. We then removed all data from the reprojected file corresponding with permanent water bodies. We note that this step will have removed permanent bodies of freshwater from our land cover map. Subsequently, this approach differentiates terrestrial and aquatic regions (henceforth, terrestrial and marine). By overlaying this terrestrial land cover map on our global complexity rasters, we were subsequently able to assign each grid cell to its corresponding terrestrial or marine classification. Using the *BayesFactor* package (*120*), we carried out Bayesian two-sample t-tests to compute Bayes Factors testing whether the sample distributions of each complexity characteristic (*x*) differ between grid cells associated with the marine and terrestrial realms (*i.e.*, *x ~ Realm*).

We observed a clear difference in the fractal dimension of marine and terrestrial environments. The distributions of *ΔH*, *R*, and *D* all differ between the marine and terrestrial realms (Fig. S2; $BF_{10} > 100$ across all pairwise comparisons); a pattern that is most evident in fractal dimension, with a greater skew towards less fractal environments in the marine environment (Fig. S2A). Although their relative distributions differ, it is intuitive that the spread of *R* and *ΔH* estimates across marine and terrestrial environments are comparable, with plains, canyons, valleys, and mountain ranges forming keystone features across global land- and seascapes. Compared to the terrestrial realm, however, the global marine environment is typically less fractal (Fig. S2A).

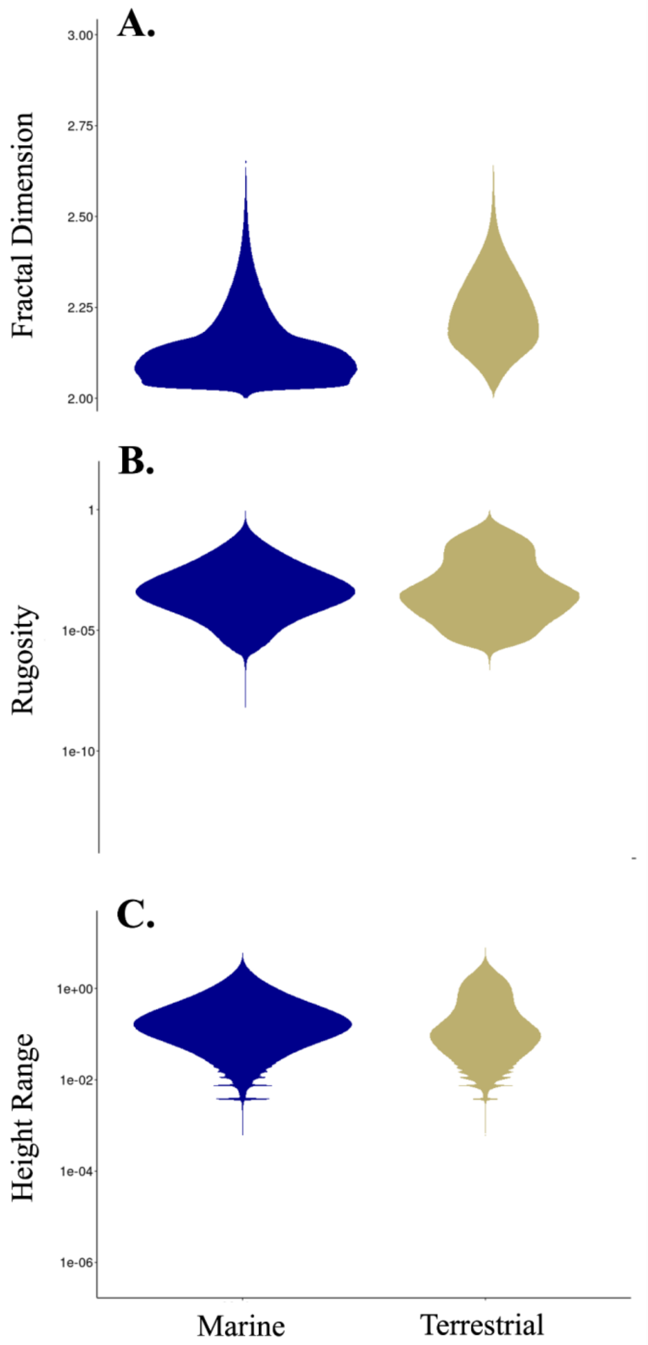


**Figure S2. Structural complexity regimes differ between marine and terrestrial environments.** The relative distributions of **(A)** fractal dimension (*D*), **(B)** rugosity (*R*), and **(C)** height range (*ΔH*) estimates across the marine and terrestrial realms.

S3. Evaluating the sensitivity of complexity measures to scale (*L*)

We tested the sensitivity of our rugosity and fractal dimension measures to changes in the scale at which they are computed by repeatedly calculating each measure for a series of successively smaller grid cells selected from the same region of our global digital elevation model (DEM). Initially, we isolated a 56.1 × 56.1 km section of our DEM (origin: x = 2,500,000, y = 0 [Mollweide equal area projection]). Iteratively working with successive values of *L* along the sequence 374, 561, 748, 935, 1122, 1870, and 18700 (all in metres), we decomposed the DEM section into a series of $L \times L$ grid cells (for $L$ = 374 this equates to 22,500 grid cells, and for $L$ = 18700 this equates to 9 grid cells). For each grid cell, we computed an estimate of its surface rugosity and fractal dimension using the corresponding functions from the *habtools* package (*98*), each time retaining these complexity estimates and the associated scale (*L*) at which they were calculated. Geometric theory dictates that rugosity and fractal dimension are additive exponents; thus, aggregating the mean estimates of these two measures from across grid cells of equal size should result in corresponding estimates for our selected area. Figure S1 illustrates the outcome of aggregating mean estimates of rugosity and fractal dimension calculated at our successive levels of L, demonstrating the consistency of our rugosity estimates across scales. Meanwhile, our estimates of fractal dimension remained largely consistent down to a scale of 935 m, after which they increased exponentially (Fig. S3). Accordingly, selecting a final scale of 1870 m for our subsequent analyses ensured that we were able to balance estimating structural complexity at a scale more intuitive for inferring ecological processes (*i.e.*, ~1km), whilst minimising the effect of resolution upon the accuracy of our complexity estimates.

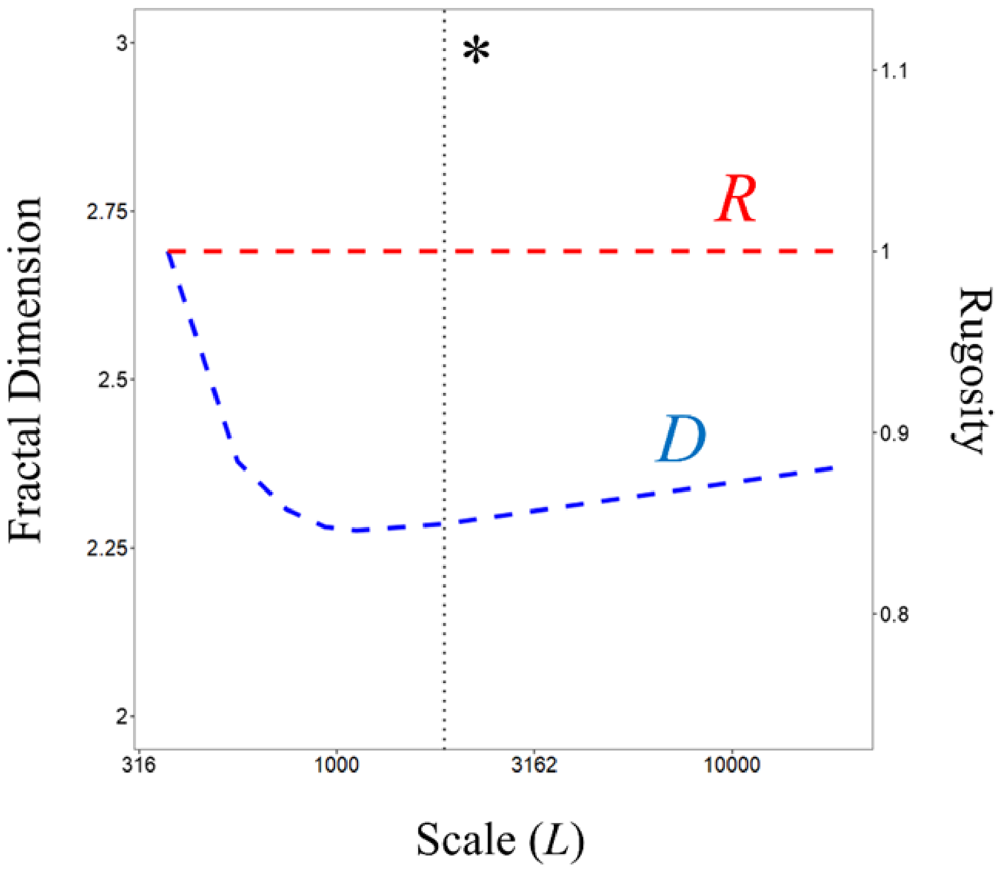


**Figure S3. The sensitivities of rugosity and fractal dimension to changing the scale (*L*) at which they are quantified.** Trends in the mean values of rugosity (*R*) and fractal dimension (*D*) estimates aggregated across a series of $L \times L$ grid cells iteratively extracted from an identical section of a digital elevation model. The dashed vertical line (*) highlights the selected value of *L* used across further analyses (1870 m).

#### S4. Selected ecosystem functional types and land cover scenarios

**Table S1. Ecosystem functional types for which estimates of rugosity and fractal dimension were computed.** Ecosystem types are shaded according to their associated realm (Freshwater, Coastal, Subterranean, Terrestrial, Wetland, or Marine). Ecosystem types correspond with the functional ecosystem groups outlined in the IUCN Global Ecosystem Typology Framework (*5*). The table continues over two pages.

| Freshwater | Coastal | Subterranean | Terrestrial | Wetlands | Marine |
|---|---|---|---|---|---|
| Permanent upland streams | River deltas | Aerobic caves | Tropical lowland rainforests | Tropical flooded peat forests | Deepwater coastal inlets |
| Permanent lowland rivers | Intertidal forests | Endolithic systems | Tropical dry forests | Permanent marshes | Riverine estuaries and bays |
| Freeze-thaw streams | Coastal saltmarsh | Underground streams and pools | Tropical montane rainforests | Seasonal floodplain marshes | Closed open inlets |
| Seasonal upland streams | Rocky shores | Groundwater ecosystems | Tropical heath forests | Episodic arid floodplains | Seagrass meadows |
| Seasonal lowland rivers | Muddy shores | Anchialine caves | Boreal montane forests | Boreal temperate bogs | Kelp forests |
| Episodic arid rivers | Sandy shores | Anchialine pools | Deciduous temperate forests | Boreal temperate fens | Photic coral reefs |
| Large rivers | Cobble shores | Sea Caves | Oceanic temperate rainforests | | Shellfish beds and reefs |
| Large permanent lakes | Coastal shrublands and grasslands | | Warm temperate rainforests | | Marine animal forests |
| Subglacial lakes | Artificial shores | | Temperate pyric humid forests | | Subtidal rock reefs |
| Small permanent lakes | | | Temperate sclerophyll forests | | Subtidal sand beds |
| Seasonal lakes | | | Seasonal dry tropical shrublands | | Subtidal mud plains |
| Freeze-thaw lakes | | | Seasonal dry temperate shrublands | | Upwelling zones |
| Ephemeral lakes | | | Cool temperate heathlands | | Epipelagic waters |
| Permanent salt lakes | | | Rocky pavements | | Mesopelagic ocean waters |
| Ephemeral salt lakes | | | Trophic savannas | | Bathypelagic ocean waters |

| | | |
|---|---|---|
| Artesian springs oases | Pyric tussock savannas | Abyssopelagic ocean waters |
| Geothermal wetlands | Hummock savannas | Sea ice |
| Large reservoirs | Temperate woodlands | Continental slopes |
| Constructed lacustrine wetlands | Temperate grasslands | Submarine canyons |
| Rice paddies | Semi-desert steppe | Abyssal plains |
| Canals and drains | Succulent Thorny deserts | Seamounts |
| | Sclerophyll hot deserts | Deepwater biogenic beds |
| | Cool temperate deserts | Hadal |
| | Hyper-arid deserts | Chemosynthetic |
| | Permanent snow | Submerged Artificial structures |
| | Polar alpine rock | Marine aquafarms |
| | Polar tundra | |
| | Temperate alpine grasslands | |
| | Trop alpine grassland | |
| | Croplands | |
| | Sown pastures and fields | |
| | Plantations | |
| | Urban and industrial | |
| | Semi-natural old fields | |

**Table S2. Land cover scenarios for which estimates of rugosity and fractal dimension were estimated.** Land cover scenarios colour coded by status (Natural [Shaded] or Anthropogenic [Unshaded]). Land cover scenarios generated using remotely sensed land cover maps sourced from the Copernicus Climate Change Service (C3S) (*35*). The table shows the number of C3S land cover sub-categories represented in each scenario, along with their associated European Space Agency Climate Change Initiative ID codes, which are shown in superscript (*121*).

| Land Cover Scenario | C3S Categories |
|---|---|
| Urban | 1 [190] |
| Cropland | 4 [10, 11, 12, 20] |
| Mosaic Vegetation/Cropland | 2 [30, 40] |
| Mosaic Vegetation | 2 [100, 110] |
| Grassland | 1 [130] |
| Shrubland | 3 [120, 121, 122] |
| Lichens & Mosses | 1 [140] |
| Sparse Vegetation | 3 [150, 152, 153] |
| Mixed Tree Cover | 1 [90] |
| Tree Cover (Needle-leaved) | 6 [70, 71, 72, 80, 81, 82] |
| Tree Cover (Broad-leaved) | 4 [50, 60, 61, 62] |
| Wetlands | 3 [160, 170, 180] |
| Bare Substrate | 3 [200, 201, 202] |
| Permanent Snow & Ice | 1 [220] |

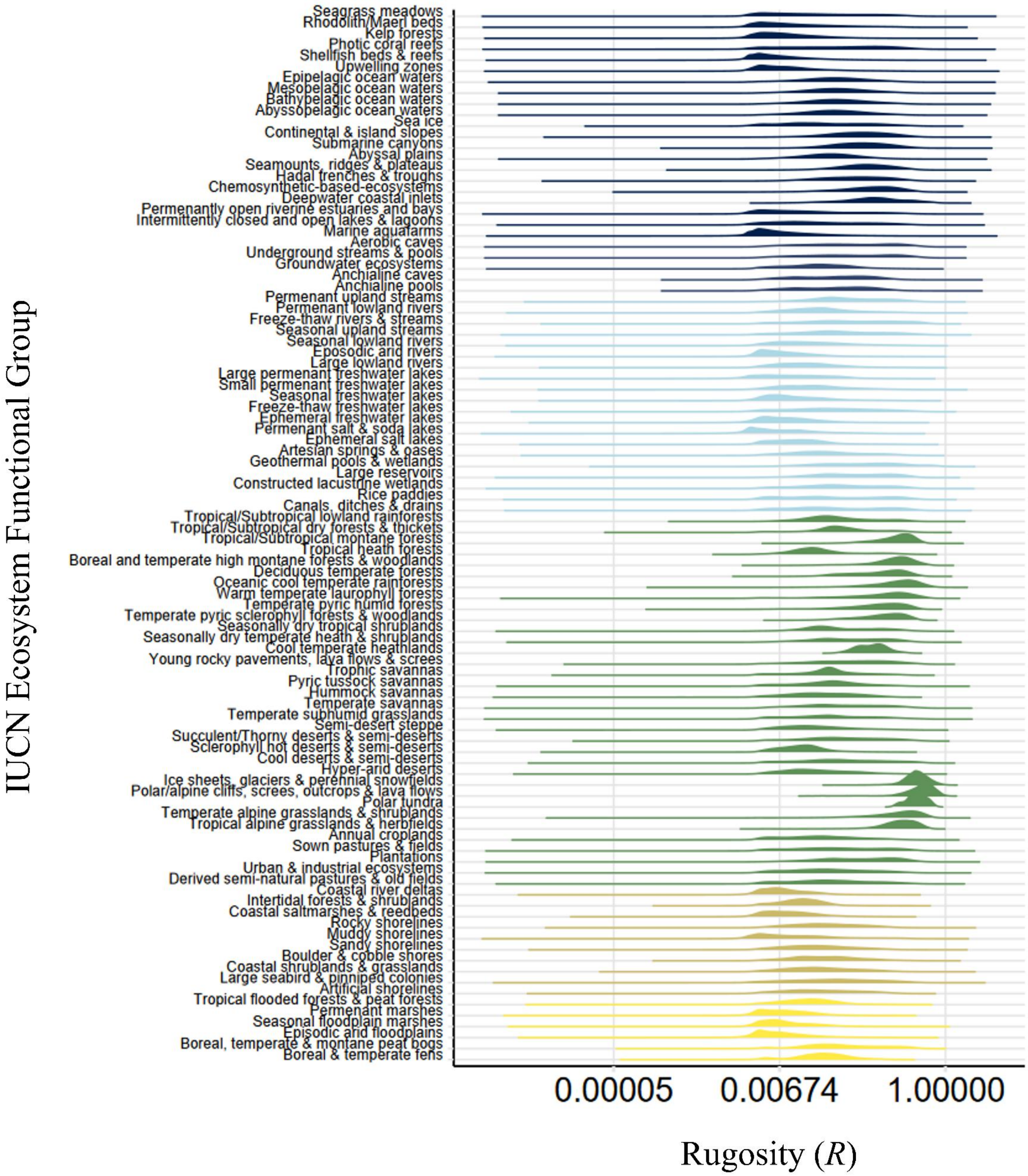
Seagrass meadows
Rhodolith/Maerl beds
Kelp forests
Photic coral reefs
Shellfish beds & reefs
Upwelling zones
Epipelagic ocean waters
Mesopelagic ocean waters
Bathypelagic ocean waters
Abyssopelagic ocean waters
Sea ice
Continental & island slopes
Submarine canyons
Abyssal plains
Seamounts, ridges & plateaus
Hadal trenches & troughs
Chemosynthetic-based-ecosystems
Deepwater coastal inlets
Permenantly open riverine estuaries and bays
Intermittently closed and open lakes & lagoons
Marine aquafarms
Aerobic caves
Underground streams & pools
Groundwater ecosystems
Anchialine caves
Anchialine pools
Permenant upland streams
Permenant lowland rivers
Freeze-thaw rivers & streams
Seasonal upland streams
Seasonal lowland rivers
Eposodic arid rivers
Large lowland rivers
Large permenant freshwater lakes
Small permenant freshwater lakes
Seasonal freshwater lakes
Freeze-thaw freshwater lakes
Ephemeral freshwater lakes
Permenant salt & soda lakes
Ephemeral salt lakes
Artesian springs & oases
Geothermal pools & wetlands
Large reservoirs
Constructed lacustrine wetlands
Rice paddies
Canals, ditches & drains
Tropical/Subtropical lowland rainforests
Tropical/Subtropical dry forests & thickets
Tropical/Subtropical montane forests
Tropical heath forests
Boreal and temperate high montane forests & woodlands
Deciduous temperate forests
Oceanic cool temperate rainforests
Warm temperate laurophyll forests
Temperate pyric humid forests
Temperate pyric sclerophyll forests & woodlands
Seasonally dry tropical shrublands
Seasonally dry temperate heath & shrublands
Cool temperate heathlands
Young rocky pavements, lava flows & screes
Trophic savannas
Pyric tussock savannas
Hummock savannas
Temperate savannas
Temperate subhumid grasslands
Semi-desert steppe
Succulent/Thorny deserts & semi-deserts
Sclerophyll hot deserts & semi-deserts
Cool deserts & semi-deserts
Hyper-arid deserts
Ice sheets, glaciers & perennial snowfields
Polar/alpine cliffs, screes, outcrops & lava flows
Polar tundra
Temperate alpine grasslands & shrublands
Tropical alpine grasslands & herbfields
Annual croplands
Sown pastures & fields
Plantations
Urban & industrial ecosystems
Derived semi-natural pastures & old fields
Coastal river deltas
Intertidal forests & shrublands
Coastal saltmarshes & reedbeds
Rocky shorelines
Muddy shorelines
Sandy shorelines
Boulder & cobble shores
Coastal shrublands & grasslands
Large seabird & pinniped colonies
Artificial shorelines
Tropical flooded forests & peat forests
Permenant marshes
Seasonal floodplain marshes
Episodic arid floodplains
Boreal, temperate & montane peat bogs
Boreal & temperate fens
IUCN Ecosystem Functional Group
0.00005
0.00674
1.00000
Rugosity (R)

**Figure S4. Comparing the rugosities of differing ecosystem functional types worldwide.** The distribution of rugosity (*R*) estimates isolated for each of the ecosystem functional groups (EFGs) identified within the IUCN's ecosystem typology (*5*) (Table S1). Ridgeline fill colour denotes the realm associated with each EFG (Marine [Navy], Freshwater [Blue], Subterranean [Purple], Terrestrial [Green], Coastal [Brown] & Wetland [Yellow]).

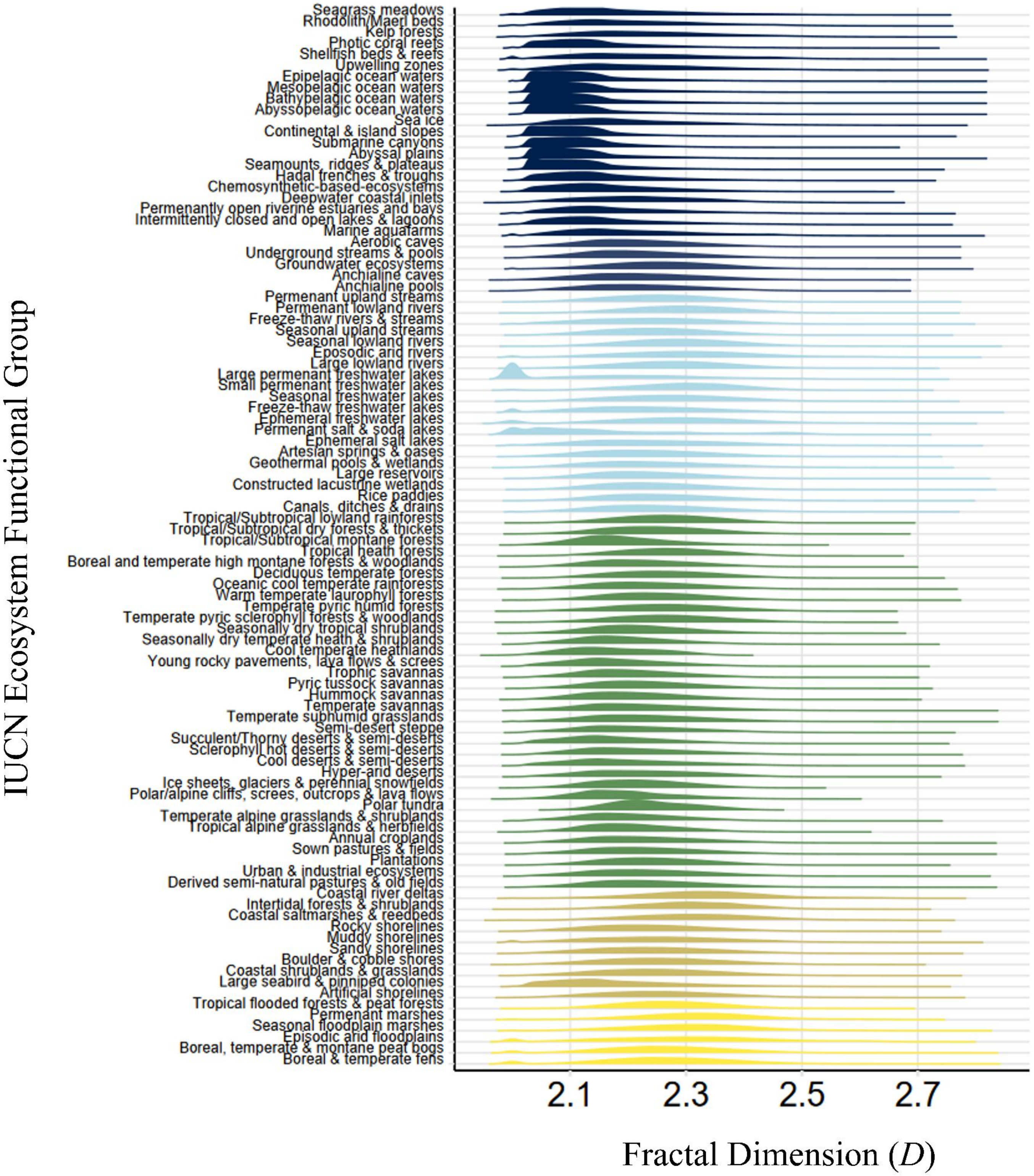
IUCN Ecosystem Functional Group
Seagrass meadows
Rhodolith/Maerl beds
Kelp forests
Photic coral reefs
Shellfish beds & reefs
Upwelling zones
Epipelagic ocean waters
Mesopelagic ocean waters
Bathypelagic ocean waters
Abyssopelagic ocean waters
Sea ice
Continental & island slopes
Submarine canyons
Abyssal plains
Seamounts, ridges & plateaus
Hadal trenches & troughs
Chemosynthetic-based-ecosystems
Deepwater coastal inlets
Permenantly open riverine estuaries and bays
Intermittently closed and open lakes & lagoons
Marine aquafarms
Aerobic caves
Underground streams & pools
Groundwater ecosystems
Anchialine caves
Anchialine pools
Permenant upland streams
Permenant lowland rivers
Freeze-thaw rivers & streams
Seasonal upland streams
Seasonal lowland rivers
Eposodic arid rivers
Large lowland rivers
Large permenant freshwater lakes
Small permenant freshwater lakes
Seasonal freshwater lakes
Freeze-thaw freshwater lakes
Ephemeral freshwater lakes
Permenant salt & soda lakes
Ephemeral salt lakes
Artesian springs & oases
Geothermal pools & wetlands
Large reservoirs
Constructed lacustrine wetlands
Rice paddies
Canals, ditches & drains
Tropical/Subtropical lowland rainforests
Tropical/Subtropical dry forests & thickets
Tropical/Subtropical montane forests
Tropical heath forests
Boreal and temperate high montane forests & woodlands
Deciduous temperate forests
Oceanic cool temperate rainforests
Warm temperate laurophyll forests
Temperate pyric humid forests
Temperate pyric sclerophyll forests & woodlands
Seasonally dry tropical shrublands
Seasonally dry temperate heath & shrublands
Cool temperate heathlands
Young rocky pavements, lava flows & screes
Trophic savannas
Pyric tussock savannas
Hummock savannas
Temperate savannas
Temperate subhumid grasslands
Semi-desert steppe
Succulent/Thorny deserts & semi-deserts
Sclerophyll hot deserts & semi-deserts
Cool deserts & semi-deserts
Hyper-arid deserts
Ice sheets, glaciers & perennial snowfields
Polar/alpine cliffs, screes, outcrops & lava flows
Polar tundra
Temperate alpine grasslands & shrublands
Tropical alpine grasslands & herbfields
Annual croplands
Sown pastures & fields
Plantations
Urban & industrial ecosystems
Derived semi-natural pastures & old fields
Coastal river deltas
Intertidal forests & shrublands
Coastal saltmarshes & reedbeds
Rocky shorelines
Muddy shorelines
Sandy shorelines
Boulder & cobble shores
Coastal shrublands & grasslands
Large seabird & pinniped colonies
Artificial shorelines
Tropical flooded forests & peat forests
Permenant marshes
Seasonal floodplain marshes
Episodic arid floodplains
Boreal, temperate & montane peat bogs
Boreal & temperate fens
2.1
2.3
2.5
2.7
Fractal Dimension (D)

**Figure S5. Comparing the fractal dimension characteristics of differing ecosystem functional types worldwide.** The distribution of fractal dimension (*D*) estimates obtained for each of the ecosystem functional groups (EFGs) identified within the IUCN's ecosystem typology (*5*) (Table S1). Ridgeline fill colour denotes the realm associated with each EFG (Marine [Navy], Freshwater [Blue], Subterranean [Purple], Terrestrial [Green], Coastal [Brown] & Wetland [Yellow]).

S5. Island details and functional richness patterns from island species-area assessment

**Table S3. A list of the islands and their associated archipelagos for which avian species and functional richness was computed.** Each island is shown along with its corresponding avian species count sourced from Blackburn *et al.* (*102*) and Matthews *et al.* (*103*), and area ($km^2$) extracted from the Global Islands database (*106*). The table continues over 16 pages.

| Archipelago | Island | Species Count | Island area ($km^2$) |
|---|---|---|---|
| Ascension | Ascension | 11 | 98.1851 |
| Azores | Corvo | 15 | 17.37523 |
| Azores | Faial | 19 | 171.7781 |
| Azores | Flores | 16 | 140.8485 |
| Azores | Graciosa | 20 | 60.40615 |
| Azores | Pico | 19 | 442.5808 |
| Azores | Sao Jorge | 20 | 244.3315 |
| Azores | Santa Maria | 20 | 97.81617 |
| Azores | Sao Miguel | 23 | 744.7828 |
| Azores | Terceira | 21 | 399.7107 |
| Bermuda | Bermuda | 11 | 40.30335 |
| Canaries | Isla de Alegranza | 18 | 10.55752 |
| Canaries | Hierro | 45 | 272.801 |
| Canaries | Fuerteventura | 45 | 1671.277 |
| Canaries | Gomera | 47 | 368.044 |
| Canaries | Graciosa Island | 20 | 28.59675 |
| Canaries | Gran Canaria | 56 | 1566.145 |
| Canaries | Lanzarote | 42 | 817.8918 |
| Canaries | La Palma | 40 | 707.7832 |
| Canaries | Lobos | 14 | 5.176882 |
| Canaries | Isla de Montaña Clara | 11 | 0.036185 |
| Canaries | Tenerife | 59 | 2047.169 |
| Cape Verdes | Boa Vista | 19 | 642.2749 |
| Cape Verdes | Ilhéu Branco | 10 | 2.993048 |

| | | | |
|---|---|---|---|
| Cape Verdes | Ilha Brava | 16 | 64.91408 |
| Cape Verdes | Fogo | 16 | 470.4217 |
| Cape Verdes | Ilha do Maio | 15 | 273.0517 |
| Cape Verdes | Ilhéu Raso | 14 | 5.775562 |
| Cape Verdes | Sal | 11 | 224.0107 |
| Cape Verdes | Santo Antao | 16 | 792.5479 |
| Cape Verdes | Santiago | 27 | 1007.154 |
| Cape Verdes | Ilha de Santa Luzia | 6 | 34.7532 |
| Cape Verdes | Sao Nicolau | 21 | 347.8258 |
| Cape Verdes | Sao Vicente | 10 | 226.0461 |
| Gulf of Guinea | Isla de Annobón | 9 | 19.94475 |
| Gulf of Guinea | Principe | 38 | 138.2783 |
| Gulf of Guinea | Sao Tome | 55 | 853.112 |
| Madeira | Madeira | 32 | 740.8338 |
| Madeira | Porto Santo | 32 | 42.23285 |
| South Atlantic | Beauchene | 16 | 1.621406 |
| South Atlantic | Bouvetøya | 11 | 48.19857 |
| South Atlantic | Gough | 22 | 64.73132 |
| South Atlantic | Inaccessible | 20 | 14.22741 |
| South Atlantic | Nightingale | 16 | 2.755008 |
| South Atlantic | South Georgia | 31 | 3561.441 |
| South Atlantic | Tristan da Cunha | 12 | 94.91268 |
| St Helena | St Helena | 10 | 123.2689 |
| Bahamas | Great Abaco | 73 | 1141.689 |
| Bahamas | North Andros | 75 | 3427.037 |
| Bahamas | Cat | 58 | 359.7128 |
| Bahamas | Eleuthera | 63 | 432.5175 |
| Bahamas | Grand Bahama | 64 | 956.1726 |
| Bahamas | Long | 58 | 407.9644 |
| Bahamas | Mayaguana | 48 | 286.8636 |
| Bahamas | New Providence | 66 | 216.4585 |

| | | | |
|---|---|---|---|
| Bahamas | San Salvador | 55 | 150.6644 |
| Caymans | Cayman Brac | 31 | 39.28784 |
| Caymans | Grand Cayman | 41 | 203.2791 |
| Caymans | Little Cayman | 30 | 29.714 |
| Greater Antilles | Cuba | 132 | 105468.3 |
| Greater Antilles | Hispaniola | 124 | 74365.9 |
| Greater Antilles | Jamaica | 100 | 11037.29 |
| Greater Antilles | Puerto Rico | 92 | 8759.471 |
| Lesser Antilles | Anguilla | 33 | 72.95279 |
| Lesser Antilles | Barbados | 34 | 435.0767 |
| Lesser Antilles | Dominica | 54 | 755.1412 |
| Lesser Antilles | Grenada | 47 | 315.6716 |
| Lesser Antilles | Guadeloupe | 57 | 1451.655 |
| Lesser Antilles | Martinique | 55 | 1113.423 |
| Lesser Antilles | Montserrat | 35 | 102.8672 |
| Lesser Antilles | Isla de Providencia | 14 | 20.85731 |
| Lesser Antilles | Saba Island | 26 | 12.97891 |
| Lesser Antilles | Isla de San Andrés | 24 | 27.58636 |
| Lesser Antilles | Île Saint-Barthélemy | 36 | 19.13918 |
| Lesser Antilles | Sint Eustatius | 26 | 20.88575 |
| Lesser Antilles | Saint Lucia | 51 | 607.2637 |
| Lesser Antilles | Saint Martin/Sint Maarten | 40 | 89.71065 |
| Antarctic | Nouvelle Amsterdam | 10 | 56.73355 |
| Antarctic | Kerguelen | 35 | 6576.191 |
| Antarctic | Marion | 27 | 296.7293 |
| Antarctic | Prince Edward | 29 | 46.18419 |
| Comoros | Anjouan | 35 | 432.8568 |
| Comoros | Grande Comore | 39 | 1018.607 |
| Comoros | Mayotte | 31 | 360.3037 |
| Comoros | Mwali | 36 | 206.3014 |
| Madagascar | Madagascar | 192 | 592521.4 |

| | | | |
|---|---|---|---|
| North Keeling | North Keeling | 11 | 2.226936 |
| Seychelles | Aride | 21 | 0.637187 |
| Seychelles | Bird | 13 | 0.940088 |
| Seychelles | Cousin | 25 | 0.324646 |
| Seychelles | Frigate | 21 | 1.964504 |
| Seychelles | La Digue | 23 | 9.818831 |
| Seychelles | Mahe | 24 | 155.5043 |
| Seychelles | Silhouette | 20 | 20.30261 |
| Mascarenes | Mauritius | 20 | 1872.954 |
| Mascarenes | Reunion | 19 | 2520.23 |
| Mascarenes | Rodrigues | 9 | 109.3016 |
| Australasia | Antipodes | 28 | 18.41531 |
| Australasia | Auckland | 34 | 461.8237 |
| Australasia | Campbell | 24 | 108.3005 |
| Australasia | Little Solander | 14 | 0.080965 |
| Australasia | Lord Howe | 18 | 16.28685 |
| Australasia | Macquarie | 26 | 128.8345 |
| Australasia | New Caledonia | 92 | 16368.26 |
| Australasia | Norfolk | 25 | 36.84829 |
| Australasia | Snares | 22 | 4.362366 |
| Australasia | Solander | 16 | 1.028442 |
| Carolines | Kosrae Island | 13 | 116.1721 |
| Carolines | Pohnpei | 28 | 351.9664 |
| Carolines | Yap Island | 19 | 92.11105 |
| Chatham Islands | Mangere | 16 | 1.153578 |
| Cook Islands | Aitutaki | 5 | 15.81133 |
| Cook Islands | Atiu | 9 | 28.816 |
| Cook Islands | Mangaia | 13 | 49.34875 |
| Cook Islands | Mauke | 5 | 19.24213 |
| Cook Islands | Mitiaro | 8 | 23.53747 |
| Cook Islands | Nassau | 2 | 1.199383 |

| | | | |
|---|---|---|---|
| Cook Islands | Rarotonga | 9 | 68.09365 |
| Cook Islands | Takutea | 8 | 1.254182 |
| Easter | Easter | 1 | 163.5291 |
| Fiji | Gau | 53 | 141.8929 |
| Fiji | Kadavu | 55 | 444.4903 |
| Fiji | Ovalau | 53 | 106.0054 |
| Fiji | Rotuma | 24 | 44.86449 |
| Fiji | Vanua Levu (West) | 61 | 283.0197 |
| Fiji | Vanua Levu (East) | 61 | 5816.859 |
| Fiji | Fiji | 62 | 10691.25 |
| Galapagos | Espanola | 22 | 61.34017 |
| Galapagos | Fernandina | 36 | 648.2214 |
| Galapagos | Santa Maria | 35 | 174.4558 |
| Galapagos | Genovesa | 21 | 14.05219 |
| Galapagos | Isabela | 42 | 4737.221 |
| Galapagos | Pinzon | 21 | 18.05281 |
| Galapagos | San Cristobal | 37 | 561.7388 |
| Galapagos | Santa Cruz | 39 | 992.8805 |
| Galapagos | Santa Fe | 22 | 24.77559 |
| Galapagos | Santiago | 38 | 578.0149 |
| Hawaiian Islands | Hawaii | 23 | 10492.91 |
| Hawaiian Islands | Kauai | 25 | 1445.693 |
| Hawaiian Islands | Lanai | 10 | 366.4668 |
| Hawaiian Islands | Maui | 15 | 1893.19 |
| Hawaiian Islands | Molokai | 13 | 679.167 |
| Hawaiian Islands | Oahu | 16 | 1559.821 |
| Kiribati | Nauru | 9 | 24.49364 |
| Line Islands | Christmas | 5 | 138.5341 |
| Marianas | Aguijan | 16 | 7.125604 |
| Marianas | Agrihan | 7 | 44.32976 |
| Marianas | Alamagan | 6 | 13.25122 |

| | | | |
|---|---|---|---|
| Marianas | Anatahan | 6 | 33.29497 |
| Marianas | Asuncion | 7 | 8.024684 |
| Marianas | Guam | 8 | 546.3411 |
| Marianas | Guguan | 14 | 4.353405 |
| Marianas | Medinilla | 8 | 0.788871 |
| Marianas | Pagan | 11 | 48.67526 |
| Marianas | Rota | 16 | 86.30281 |
| Marianas | Saipan | 17 | 120.8604 |
| Marianas | Sarigan | 6 | 4.437911 |
| Marianas | Tinian | 17 | 102.5831 |
| Marianas | Farallon de Pajaros | 6 | 2.30647 |
| Marquesas | Eiao | 3 | 36.00569 |
| Marquesas | Fatu Hiva | 7 | 83.77272 |
| Marquesas | Fatu Huku | 1 | 0.569641 |
| Marquesas | Hatutaa | 4 | 6.793498 |
| Marquesas | Hiva Oa | 5 | 306.0223 |
| Marquesas | Motane | 4 | 12.74041 |
| Marquesas | Nuku Hiva | 8 | 337.008 |
| Marquesas | Tahuata | 5 | 68.09724 |
| Marquesas | Ua Huka | 4 | 86.3498 |
| Marquesas | Ua Pou | 5 | 103.5066 |
| Pitcairns | Henderson | 14 | 44.35862 |
| Pitcairns | Oeno | 12 | 0.756411 |
| Samoa | Aunuu | 21 | 1.484449 |
| Samoa | Ofu | 23 | 7.397082 |
| Samoa | Olosega | 23 | 5.786605 |
| Samoa | Maafee | 11 | 0.059253 |
| Samoa | Savaii | 40 | 1714.469 |
| Samoa | Swains | 10 | 3.912446 |
| Samoa | Ta’u | 29 | 47.0725 |
| Samoa | Tutuila | 27 | 137.2652 |

| | | | |
|---|---|---|---|
| Samoa | Upolu | 38 | 1133.162 |
| Society Islands | Mai'ao | 7 | 12.73089 |
| Society Islands | Meheti'a | 4 | 2.107403 |
| Society Islands | Moorea | 11 | 133.7528 |
| Society Islands | Tahiti | 15 | 1051.288 |
| Tonga | Eua | 25 | 87.30712 |
| Tonga | Niuafo'ou | 15 | 51.73785 |
| Tonga | Niue | 15 | 262.983 |
| Tuamotus | Makatea | 11 | 29.59867 |
| Tubuai | Rapa | 21 | 38.53623 |
| Vanuatu | Aneityum | 31 | 160.7907 |
| Vanuatu | Efate | 43 | 898.4524 |
| Vanuatu | Erromango | 36 | 892.3798 |
| Vanuatu | Espiritu Santo | 49 | 3959.578 |
| Vanuatu | Malakula | 43 | 2058.196 |
| Vanuatu | Tanna | 33 | 563.3906 |
| Dahlak | Nocra | 12 | 6.058826 |
| Dahlak | Seil Nocra | 5 | 0.186948 |
| Dahlak | Entedebir | 7 | 1.531689 |
| Dahlak | Enteraya | 4 | 1.323061 |
| Dahlak | Dur-Ghella | 3 | 0.324737 |
| Dahlak | Dur Gaam | 2 | 0.277853 |
| Dahlak | Sarad | 5 | 1.244417 |
| Dahlak | Dar Ottun | 6 | 2.219403 |
| Dahlak | Duliacus | 5 | 0.463875 |
| Dahlak | Dalcus | 4 | 0.360378 |
| Dahlak | Shumma | 9 | 5.745295 |
| Dahlak | Madote | 2 | 0.040468 |
| Dahlak | Ota | 2 | 1.12499 |
| Dahlak | Dissei | 17 | 7.025622 |
| Dahlak | Sheik Said | 5 | 0.406695 |

| | | | |
|---|---|---|---|
| Dahlak | Harat | 7 | 23.05943 |
| Dahlak | Sheik el Abu | 2 | 0.22715 |
| Dahlak | Dehil | 7 | 12.58565 |
| Dahlak | Baradu | 5 | 2.844789 |
| Dahlak | Dehil Bahut | 4 | 0.526908 |
| Dahlak | Dahret | 2 | 0.339448 |
| Faroe Islands | Suduroy | 41 | 177.0059 |
| Faroe Islands | Litla Dimun | 7 | 0.963461 |
| Faroe Islands | Stóra Dímun | 13 | 121.421 |
| Faroe Islands | Skúvoy | 20 | 11.41223 |
| Faroe Islands | Sandoy | 41 | 117.3093 |
| Faroe Islands | Hestur | 21 | 7.487726 |
| Faroe Islands | Koltur | 13 | 3.006029 |
| Faroe Islands | Mykines | 25 | 12.12271 |
| Faroe Islands | Tindhólmur | 9 | 1.383596 |
| Faroe Islands | Vagar | 31 | 180.324 |
| Faroe Islands | Nólsoy | 27 | 10.95951 |
| Faroe Islands | Streymoy | 38 | 381.0073 |
| Faroe Islands | Eysturoy | 38 | 288.6772 |
| Faroe Islands | Kalsoy | 20 | 31.71933 |
| Faroe Islands | Kunoy | 21 | 35.64738 |
| Faroe Islands | Bordoy | 51 | 137.8889 |
| Faroe Islands | Svínoy | 19 | 27.27977 |
| Faroe Islands | Fugloy | 21 | 10.99905 |
| Ryukyus | Hateruma-jima | 31 | 12.93636 |
| Ryukyus | Nakanokami-shima | 21 | 0.220247 |
| Ryukyus | Kuro-shima | 30 | 10.23009 |
| Ryukyus | Taketomi-shima | 23 | 5.516685 |
| Ryukyus | Obama-jima | 43 | 7.960337 |
| Ryukyus | Iriomote-jima | 59 | 290.8561 |
| Ryukyus | Yonaguni-jima | 45 | 28.50496 |

| | | | |
|---|---|---|---|
| Ryukyus | Rasa-jima | 15 | 1.160107 |
| Ryukyus | Ishigaki Jima | 57 | 225.6296 |
| Ryukyus | Hatoma-jima | 16 | 0.987339 |
| Ryukyus | Tarama-jima | 38 | 20.09645 |
| Ryukyus | Kurima-jima | 39 | 2.943458 |
| Ryukyus | Minna-shima | 11 | 2.248897 |
| Ryukyus | Miyako | 49 | 161.7133 |
| Ryukyus | Irabu-jima | 32 | 39.83729 |
| Ryukyus | Ikema-jima | 29 | 3.049565 |
| Ryukyus | Nan Xiaodao | 15 | 0.457542 |
| Ryukyus | Bei Xiaodao | 12 | 0.351008 |
| Ryukyus | Minamidaito-jima | 36 | 30.4525 |
| Ryukyus | Taisyo-jima | 10 | 0.070914 |
| Ryukyus | Kuba-shima | 8 | 1.961366 |
| Ryukyus | Kitadaito-jima | 30 | 11.75674 |
| Ryukyus | Geruma-jima | 19 | 1.095667 |
| Ryukyus | Tokashiki-jima | 23 | 14.76232 |
| Ryukyus | Aka-shima | 24 | 3.520193 |
| Ryukyus | Yakabi-jima | 17 | 1.212998 |
| Ryukyus | Zamami-jima | 27 | 6.026995 |
| Ryukyus | Nagannu-jima | 4 | 0.161408 |
| Ryukyus | Hamahiga-jima | 16 | 2.076744 |
| Ryukyus | Kume-jima | 36 | 60.71395 |
| Ryukyus | Henza-jima | 25 | 10.58901 |
| Ryukyus | Tonaki-jima | 10 | 3.713011 |
| Ryukyus | Ikei-jima | 12 | 1.639206 |
| Ryukyus | Okinawa-jima | 55 | 1209.311 |
| Ryukyus | Aguni-jima | 41 | 7.792755 |
| Ryukyus | Sesoko-jima | 16 | 2.87777 |
| Ryukyus | Minna-shima | 12 | 0.453393 |
| Ryukyus | Yagaji-shima | 25 | 7.044524 |

| | | | |
|---|---|---|---|
| Ryukyus | Kouri-jima | 18 | 2.753749 |
| Ryukyus | Ie-jima | 25 | 21.06218 |
| Ryukyus | Izena-jima | 28 | 13.86452 |
| Ryukyus | Iheya-jima | 34 | 20.38432 |
| Ryukyus | Yoron-jima | 40 | 20.28754 |
| Ryukyus | Okinoerabu-jima | 40 | 94.17043 |
| Ryukyus | Toku-no-shima | 50 | 252.9811 |
| Ryukyus | Uke-shima | 34 | 13.17055 |
| Ryukyus | Yoro-shima | 36 | 9.17498 |
| Ryukyus | Kakeroma-jima | 44 | 77.61778 |
| Ryukyus | Amami-O-shima | 65 | 712.9028 |
| Ryukyus | Kikai-jima | 38 | 56.26759 |
| Ryukyus | Takara-jima | 32 | 7.307032 |
| Ryukyus | Kodakara-jima | 14 | 0.990809 |
| Ryukyus | Akuseki-to | 33 | 7.039042 |
| Ryukyus | Suwanose-jima | 29 | 26.28518 |
| Ryukyus | Taira-jima | 52 | 2.045565 |
| Ryukyus | Naka-no-shima | 55 | 34.22619 |
| Ryukyus | Gajya-jima | 24 | 4.080054 |
| Ryukyus | Kuchi-no-shima | 28 | 13.06442 |
| Ryukyus | Yaku-shima | 65 | 503.1764 |
| Ryukyus | Kuchinoerabu-jima | 13 | 35.05186 |
| Ryukyus | Tane-ga-shima | 42 | 447.0383 |
| Ryukyus | Mage-shima | 15 | 8.405635 |
| Ryukyus | Kamino-shima | 36 | 0.190987 |
| Ryukyus | Uji-mukae-jima | 12 | 1.68443 |
| Ryukyus | Uchi-jima | 28 | 0.528605 |
| New Zealand | North Island | 67 | 114573.1 |
| New Zealand | South Island | 80 | 150683.1 |
| New Zealand | Stewart | 52 | 1705.122 |
| New Zealand | Great Barrier | 38 | 286.3621 |

| | | | |
|---|---|---|---|
| New Zealand | D'Urville | 38 | 169.8388 |
| New Zealand | Little barrier | 36 | 31.39119 |
| New Zealand | Great Mercury | 27 | 20.03392 |
| New Zealand | Kapiti | 39 | 19.73914 |
| New Zealand | Codfish | 35 | 15.6124 |
| New Zealand | Mayor | 23 | 13.66832 |
| New Zealand | Big South Cape | 29 | 10.45092 |
| New Zealand | Taranga | 27 | 5.38155 |
| New Zealand | Cavalli | 22 | 4.242278 |
| New Zealand | Manawatawhi | 21 | 3.922256 |
| New Zealand | Nukuwoiota | 30 | 2.024218 |
| New Zealand | Whatupuke | 22 | 1.058761 |
| New Zealand | Tawhiti Rahi | 18 | 1.779046 |
| New Zealand | Cuvier | 18 | 2.073943 |
| New Zealand | Stephens | 33 | 1.479828 |
| New Zealand | Solander | 21 | 1.028442 |
| New Zealand | Moutohora | 26 | 1.890445 |
| Wakatobi | Wangi-wangi | 53 | 158.7033 |
| Wakatobi | Pulau Oroho | 39 | 13.19836 |
| Wakatobi | Pulau Kapota | 26 | 18.46125 |
| Wakatobi | Pulau Hoga | 35 | 3.621684 |
| Wakatobi | Pulau Kaledupa | 61 | 65.79598 |
| Wakatobi | Tomia | 44 | 53.47735 |
| Wakatobi | Pulau Lenteaoge | 14 | 16.97737 |
| Wakatobi | Pulau Binongko | 41 | 99.75983 |
| Wakatobi | Pulau Runduma | 22 | 5.486015 |
| California | San Miguel | 10 | 38.68636 |
| California | Santa Rosa | 22 | 216.2562 |
| California | Santa Cruz | 32 | 250.7135 |
| California | San Nicolas | 10 | 58.92588 |
| California | Santa Barbara | 13 | 2.59091 |

| | | | |
|---|---|---|---|
| California | Santa Catalina | 28 | 194.5769 |
| California | San Clemente | 23 | 147.228 |
| California | San Martin | 4 | 2.691476 |
| California | Guadelloupe | 12 | 245.9403 |
| California | Cedros | 12 | 350.4197 |
| California | Natividad | 5 | 7.782681 |
| Haida Gwaii | Murchinson | 18 | 4.552043 |
| Haida Gwaii | Reef | 27 | 2.279018 |
| Haida Gwaii | East Limestone | 20 | 0.536084 |
| Haida Gwaii | House Island | 16 | 0.344765 |
| Haida Gwaii | De la Beche Island | 12 | 0.290071 |
| Haida Gwaii | Hotspring Island | 19 | 0.196239 |
| Haida Gwaii | Agglomerate Island | 18 | 0.237544 |
| Haida Gwaii | Marco Island | 12 | 0.305898 |
| Haida Gwaii | West Limestone | 13 | 0.111418 |
| Haida Gwaii | Haswell Island | 13 | 0.131049 |
| Haida Gwaii | Helmet Island | 10 | 0.11481 |
| Haida Gwaii | Hutton Island | 8 | 0.127501 |
| Haida Gwaii | Low Island | 9 | 0.089824 |
| Haida Gwaii | Kawas Islets | 11 | 0.047127 |
| Haida Gwaii | Titul Island | 9 | 0.086687 |
| Haida Gwaii | Sivart Island | 12 | 0.082664 |
| Haida Gwaii | Faraday | 6 | 3.174856 |
| Haida Gwaii | Marco Island | 6 | 0.305898 |
| Maddalena | Isola Piana | 1 | 0.050858 |
| Maddalena | Isola Barrettini | 2 | 0.126197 |
| Maddalena | Isolotto Spargiotto | 3 | 0.127032 |
| Maddalena | Isola Corcelli | 2 | 0.143295 |
| Maddalena | Isola Razzoli | 12 | 1.813203 |
| Maddalena | Isola Santa Maria | 17 | 2.349299 |
| Maddalena | Isola Spargi | 20 | 4.388439 |

| | | | |
|---|---|---|---|
| Maddalena | Isola Maddalena | 36 | 20.95371 |
| Aegean | Thasos | 57 | 384.7024 |
| Aegean | Samothraki | 43 | 180.8276 |
| Aegean | Lemnos | 41 | 479.3114 |
| Aegean | Nisí Ágios Efstrátios | 25 | 41.8989 |
| Aegean | Lesbos | 81 | 1637.989 |
| Aegean | Skyros | 29 | 210.5662 |
| Aegean | Khios | 43 | 846.8542 |
| Aegean | Nisí Psará | 13 | 39.00953 |
| Aegean | Nisí Antípsara | 12 | 4.265915 |
| Aegean | Samos | 60 | 478.575 |
| Aegean | Icaria | 32 | 255.6389 |
| Aegean | Nisí Foúrnoi | 18 | 31.54456 |
| Aegean | Kythira | 29 | 278.7307 |
| Aegean | Nisí Astypálaia | 21 | 98.30458 |
| Aegean | Andros | 28 | 382.1811 |
| Aegean | Tinos | 26 | 196.6755 |
| Aegean | Nisí Sýros | 22 | 85.41883 |
| Aegean | Mykonos | 18 | 87.8033 |
| Aegean | Kea | 25 | 133.0529 |
| Aegean | Nisí Kýthnos | 22 | 100.2475 |
| Aegean | Nisí Sérifos | 23 | 73.68975 |
| Aegean | Nisí Sífnos | 23 | 77.34559 |
| Aegean | Milos | 29 | 159.9123 |
| Aegean | Paros | 33 | 197.8017 |
| Aegean | Naxos | 35 | 431.475 |
| Aegean | Amorgos | 23 | 121.421 |
| Aegean | Ios | 24 | 110.0742 |
| Aegean | Nisí Síkinos | 24 | 41.93784 |
| Aegean | Nisí Folégandros | 24 | 32.67019 |
| Aegean | Santorini Island | 23 | 76.42349 |

| | | | |
|---|---|---|---|
| Aegean | Nisí Anáfi | 19 | 38.75907 |
| Aegean | Nisída Antímilos | 11 | 8.752003 |
| Aegean | Nisí Polýaigos | 12 | 18.23104 |
| Aegean | Nisí Antíparos | 25 | 35.25457 |
| Aegean | Nisí Donoúsa | 12 | 13.82711 |
| Aegean | Nisí Irákleia | 12 | 18.5268 |
| Aegean | Koufonísi | 12 | 5.923634 |
| Aegean | Nisí Schoinoússa | 12 | 9.104391 |
| Aegean | Nisída Káto Koufonísi | 12 | 4.089351 |
| Aegean | Nisída Kéros | 12 | 15.53981 |
| Aegean | Nisí Antikýthira | 8 | 20.77699 |
| Aegean | Nisí Leipsoí | 25 | 16.4139 |
| Aegean | Léros Island | 19 | 55.34684 |
| Aegean | Kalymnos Island | 18 | 112.166 |
| Aegean | Nisída Télendos | 17 | 4.787799 |
| Aegean | Nisída Kalólimnos | 25 | 2.07572 |
| Aegean | Nisída Kínaros | 15 | 4.548975 |
| Aegean | Nisída Levítha | 15 | 9.329495 |
| Aegean | Nisí Pátmos | 16 | 34.7266 |
| Aegean | Kos | 42 | 289.3308 |
| Aegean | Nisí Nísyros | 26 | 41.29873 |
| Aegean | Nisí Tílos | 25 | 62.50285 |
| Aegean | Nisí Sými | 23 | 58.99019 |
| Aegean | Nisí Chálki | 20 | 28.1828 |
| Aegean | Rhodes | 39 | 1405.939 |
| Aegean | Nisída Sariá | 13 | 20.81999 |
| Aegean | Karpathos | 18 | 303.124 |
| Aegean | Kasos Island | 18 | 66.59687 |
| Aegean | Crete | 66 | 8296.38 |
| Aegean | Nisída Ágria Gramvoúsa | 3 | 0.871926 |

| | | | |
|---|---|---|---|
| Aegean | Nisída Ímeri Gramvoúsa | 5 | 0.740241 |
| Aegean | Nisída Pontikonísi | 2 | 0.262213 |
| Aegean | Nisída Elafónisos | 3 | 0.501205 |
| Aegean | Nisída Kolokythás | 3 | 0.126725 |
| Aegean | Nisí Chrysí | 2 | 5.146783 |
| Aegean | Nisída Ágioi Theódoroi | 4 | 0.813568 |
| Aegean | Nisí Gávdos | 10 | 33.44813 |
| Aegean | Nisída Gavdopoúla | 5 | 1.840591 |
| Aegean | Nisída Paximádi | 3 | 0.02808 |
| Aegean | Nisí Día | 5 | 12.11412 |
| Aegean | Nisída Elása | 2 | 1.818937 |
| Aegean | Nisída Dragonáda | 6 | 2.875214 |
| Aegean | Nisída Gianysáda | 6 | 2.141819 |
| Aegean | Nisída Paximáda | 6 | 0.285417 |
| Aegean | Kefali | 2 | 0.007762 |
| Aegean | Nisída Kaválos | 2 | 0.022159 |
| Aegean | Koufonísi | 4 | 4.433834 |
| Riau Lingaa | Pulau Akka | 15 | 0.135325 |
| Riau Lingaa | Batam | 80 | 393.6485 |
| Riau Lingaa | Bintan | 114 | 1168.489 |
| Riau Lingaa | Pulau Durian Besar | 48 | 19.47019 |
| Riau Lingaa | Lingga | 115 | 863.5657 |
| Riau Lingaa | Pulau Malang | 10 | 0.027408 |
| Riau Lingaa | Pulau Cuma | 6 | 0.034899 |
| Riau Lingaa | Pulau Momoi | 23 | 1.304972 |
| Riau Lingaa | Pulau Nginang | 32 | 7.259278 |
| Riau Lingaa | Pulau Nibung | 7 | 0.245556 |
| Riau Lingaa | Pulau Paloi | 10 | 0.031843 |
| Riau Lingaa | Pulau Reman | 13 | 0.107739 |
| Riau Lingaa | Pulau Sambu | 12 | 0.605948 |

| | | | | |
|---|---|---|---|---|
| 1152 | Riau Lingaa | Pulau Saya | 8 | 0.888522 |
| | Riau Lingaa | Pulau Sayap | 6 | 0.005196 |
| | Riau Lingaa | Pulau Sebangka | 41 | 134.8664 |
| | Riau Lingaa | Pulau Sepatu | 11 | 0.019593 |
| | Riau Lingaa | Pulau Temiang | 41 | 41.20528 |
| | West Sumatra | Babi | 28 | 48.66654 |
| | West Sumatra | Pulau Bangkaru | 25 | 60.33863 |
| | West Sumatra | Enggano | 40 | 407.41 |
| | West Sumatra | Pulau Lasia | 13 | 15.55406 |
| | West Sumatra | Nias | 119 | 4150.049 |
| | West Sumatra | Pagai Utara | 41 | 606.3252 |
| | West Sumatra | Pini | 31 | 323.159 |
| | West Sumatra | Siberut | 93 | 3847.75 |
| | West Sumatra | Simeulue | 76 | 1750.022 |
| | West Sumatra | Sipura | 58 | 600.4683 |
| | West Sumatra | Pulau Pagai Selatan | 42 | 868.5012 |
| | West Sumatra | Tanahbala | 20 | 449.5015 |
| | West Sumatra | Tanahmasa | 51 | 338.2108 |
| | West Sumatra | Pulau Telo | 40 | 11.19593 |
| | West Sumatra | Tuangku | 54 | 205.247 |
